# Integration of a neuronal RNAseq dataset with the draft *Gryllus bimaculatus* transcriptome refines gene predictions and highlights potential systematic response to injury

**DOI:** 10.1101/2025.07.13.663756

**Authors:** Felicia F. Wang, Harrison P. Fisher, Lisa M. Ledwidge, Joel H. Graber, Riley A. Grindle, Jarod A. Rollins, Hadley W. Horch

## Abstract

The cricket *Gryllus bimaculatus* presents a compelling model for investigating neuroplasticity due to its unusual capability of adult structural reorganization. The molecular pathways underlying these changes are entirely unknown. Here, we reanalyzed RNAseq data, drawn from deafferented neuronal tissue one, three, and seven days post-injury, that was previously used to assemble a *de novo* transcriptome. In this current analysis, we aligned our original RNAseq data to the publicly available *G. bimaculatus* draft genome, and used the resulting alignments to refine and update the existing annotations. We identified over 10,000 missing genes and reported a measurable improvement in BUSCO scores. These updated annotations were then used as the basis for a DESeq2 differential expression analysis and subsequent functional enrichment analysis to further explore the potential molecular basis of this compensatory anatomical plasticity. Days one and three showed the largest post-deafferentation expression differences. Overall, more transcripts were upregulated rather than downregulated. Protein-protein interactions enriched for G-protein-related signaling, hormone metabolism, and membrane dynamics were evident. Changes in expression of factors related to small GTPases and nervous system development were particularly intriguing. We also identified a surprising enrichment of GO terms related to muscle contraction in this neuronal-specific transcriptome. Identifying these and other differentially regulated transcripts can be used to design hypotheses around well-conserved molecular mechanisms that may be involved in this unique example of adult structural plasticity in the cricket.

## Background

Most adult organisms, especially mammals, are limited in their capacity to recover from neurological damage (Prigge and Kay, 2018; Sampaio-Baptista et al., 2018). The Mediterranean field cricket, *Gryllus bimaculatus*, provides a model of neuroplasticity due to its demonstrated ability to compensate for neuronal damage with novel dendritic growth and synapse formation, even into adulthood. Specifically, the central auditory system, much of which resides in the prothoracic ganglion, reorganizes in response to deafferentation caused by unilateral transection of auditory afferents in the adult (Brodfuehrer and Hoy, 1988; Horch et al., 2011)

In crickets, auditory information is transduced by the auditory organs, located on the prothoracic limbs. Auditory afferents receive the sensory stimuli and convey this information into the prothoracic ganglion where they form synapses with several different auditory neurons (Popov et al., 1978; Poulet and Hedwig, 2001). These neurons exist as mirror image pairs and their dendritic arbors remain localized ipsilateral to the auditory input, typically not projecting contralaterally across the midline (Wohlers and Huber, 1985). However, previous research has shown that after amputation of the prothoracic leg, which removes the auditory organ and severs the afferents, the deafferented dendrites of the ipsilateral auditory neurons sprout across the midline and form functional synapses with the intact auditory afferents on the contralateral side. This reorganization is evident whether deafferentation occurs in larvae (Hoy et al., 1985; Schildberger et al., 1986) or adults (Brodfuehrer and Hoy, 1988; Horch et al., 2011; Schmitz, 1989). Various aspects of the physiological consequences of this compensatory behavior have been studied (Brodfuehrer and Hoy, 1988; Hoy et al., 1985; Schildberger et al., 1986), however little is known about the molecular pathways and mechanisms underlying this growth.

Various *de novo* transcriptomes have been created for use in *Gryllus bimaculatus* (Bando et al., 2013; Fisher et al., 2018; Zeng et al., 2013, p. 201; Zeng and Extavour, 2012), including one built with RNA from individual prothoracic ganglia of both control and deafferented adult male crickets (Fisher et al., 2018). Initially, this *de novo* transcriptome assembly was mined for the presence of developmental guidance molecules, though no differential analysis was completed. While guidance molecules have mainly been studied for their role in development, it has also been suggested that alterations in their expression may influence the ability of axons and dendrites to recover from injury in adulthood (Fisher et al., 2018; Harel and Strittmatter, 2006; Kaneko et al., 2006; Yu et al., 1998). Mining this cricket transcriptome revealed that many well-conserved developmental guidance molecules, including *slit*, *ephrins*, *netrins*, and *semaphorins*, were present within the adult prothoracic transcriptome (Fisher et al., 2018). The goal of the current study was to complete a differential analysis to better generate hypotheses focused on the underlying molecular control of the compensatory dendritic growth observed in the cricket auditory system. In this updated analysis, we aligned the cricket prothoracic ganglion RNA-seq reads (Fisher et al., 2018) to the publicly available *Gryllus bimaculatus* genome (Ylla et al., 2021a), integrated these alignments into the reference transcriptome, and finally used the resulting updated transcriptome to quantify expression across these experimental samples. We then analyzed changes in transcript expression levels one, three, and seven days post-deafferentation. The identified genes were then analyzed using gene ontology (GO) annotation analysis and protein-protein interaction enrichment analysis, to determine the types of transcripts that were differentially regulated over the course of the injury response. By performing this analysis, we discovered gene expression changes evident over the course of the compensatory growth response, allowing for the development of future hypotheses focused on pathways or key molecules critical to this process.

## Results and Discussion

### Transcriptome Assembly and Analysis

This transcriptomic study focused on the cricket, *Gryllus bimaculatus*, whose nervous system has been shown to have an unusual level of adult structural plasticity (Brodfuehrer and Hoy, 1988; Horch et al., 2011; Schmitz, 1989). We deafferented central sensory neurons, including the auditory neurons, in the prothoracic ganglion of the adult cricket by unilateral amputation of the prothoracic leg at the femur. The auditory organ resides just distal to the tibial-femoral joint on the prothoracic leg. Control amputations, designed to control for the stress of injury, consisted of removal of the distal tip of the tarsus. We harvested prothoracic ganglia one-, three-, and seven-days post-amputation. These time points were designed to capture transcriptional changes in response to the loss of activity (day one), during initial sprouting (days one and three), growth across the midline (days three and seven), and novel synapse formation (days three and seven; Brodfuehrer and Hoy, 1988; Pfister et al., 2013). Although a *de novo* assembly from *G. bimaculatus* prothoracic ganglion was completed previously (Fisher et al., 2018), the present study aligned the sequence reads to the published genome (Ylla et al., 2021a), generating updated transcriptome annotations, which were then used for differential expression analysis.

This genome-based analysis yielded 74,090 predicted transcripts from 28,637 genes (Supplemental files S1 (.gtf) and and S2 (.fa)), which was far lower than the number predicted in our *de novo* assembly (374,383 transcripts; Fisher et al. 2018). This updated transcriptome also represents a significant increase in the number of annotations over the the original draft genome assembly (Ylla et al., 2021b), which represented 28,529 transcripts from 17,871 genes (Table 1). Also, in comparison with the original genome assembly, the average and median transcript length increased from 2,624 and 1,848 nucleotides to 3,084 and 2,085 nucleotides, respectively, and the maximum transcript length increased from 27,129 to 63,870 nucleotides (Table 1).

**Table 1:** Summative detail from the original genome assembly (GBI) compared to our current, updated assembly (GBIG).

|  | GBI | GBIG<br>(update) |
| --- | --- | --- |
| N(Genes) | 17,871 | 28,637 |
| N(Transcripts) | 28,529 | 74,090 |
| Average transcripts per gene | 1.6 | 2.6 |
| Max transcripts per gene | 19 | 40 |
| Average transcript length | 2,624 | 3,084 |
| Median transcript length | 1,848 | 2,085 |
| Max transcript length | 27,129 | 63,870 |

In cases where our assembly suggested new transcript isoforms to existing GBI-annotated genes, we preserved the GBI name and annotation, but added the isoforms to the gene definition. In total, 25,438 new transcript isoforms were added to 9,301 genes. The maximum number of added transcripts was 22, added to GBI_11181, which was annotated as “Similar to Dscam2: Down syndrome cell adhesion molecule-like protein Dscam2 (Drosophila melanogaster).” The Dscam family of genes is well-known for its high number of splicing isoforms (Kerwin et al., 2018; Li and Millard, 2019; Schmucker et al., 2000).

Our updated assembly also resulted in 1,409 instances where the StringTie evidence suggested that neighboring regions on the draft genome that were originally annotated as separate genes are instead separate components of a single transcription unit. In such cases, our algorithm gave a new GBIG identifier to the complete gene, while retaining the transcript identifiers from the existing subunits from the GBI assembly. Table 2 shows the distribution of the number of genes joined together by our evidence, while Supplemental Table 1 (S3 Table) lists all such instances, including both the new identifier along with the GBI genes that were joined.

**Table 2:** Evidence-based merging of GBI gene neighbors. The revised GBIG transcriptome included multiple instances of evidence that neighboring GBI-annotated genes form a single transcriptional unit. Combinations ranged from two neighbor regions, which occurred 1,082 times, up to 11 neighbor regions, which occurred once.

| # of joined GBI neighbors | # of GBIG Instances |
| --- | --- |
| 2 | 1082 |
| 3 | 233 |
| 4 | 53 |
| 5 | 27 |
| 6 | 9 |
| 7 | 4 |
| 11 | 1 |

Finally, the updated transcriptome includes 12,587 completely novel genes, supported by a Supplemental Figure (S4), which presents several comparisons of the GBI and GBIG annotations. These Integrated Genome Viewer (IGV) representations of both annotations, as well as the reduced BAM alignment file, show the evidence supporting the updated annotations.

BUSCO (Benchmarking Universal Single-Copy Orthologs) analysis (Manni et al., 2021b, 2021a; Simão et al., 2015) is a standard approach for assessing the completeness of a genome or transcriptome. Using evolutionarily informed expectations of gene content, BUSCO analysis assesses the presence and multiplicity of genes that have been identified to be near-universally present as single-copy across defined phylogenetic ranges. The BUSCO scores for our updated transcriptome indicate a high quality assembly (Table 3), which improved upon the reference assembly (Ylla et al., 2021a). Using BUSCO version 5.7.0 and the arthropoda_odb10 reference, the number of missing genes in the updated transcriptome was reduced from 26 to 7 and the number of fragmented genes from 22 to 9.

**Table 3:** BUSCO scores for the current assembly as compared to the annotations for the recent assembly of the genome.

|  | Genome Annotation |  | Current Assembly |  |
| --- | --- | --- | --- | --- |
| **Arthropoda:** | **Count** | **%** | **Count** | **%** |
| complete- total | 965 | 95.26 | 997 | 98.42 |
| complete-single | 916 | 90.42 | 939 | 92.69 |
| complete-duplicated | 49 | 4.84 | 58 | 5.73 |
| fragmented | 22 | 2.17 | 9 | 0.89 |
| missing | 26 | 2.57 | 7 | 0.69 |
| total | 1013 |  | 1013 |  |

It is noteworthy that in the case of five BUSCO genes that were missing in the GBI assembly but present in our updated assembly, inspection showed that the newly identified matches were to genes that were present in the GBI transcriptome, however, the BUSCO matches were only identified with the addition of transcript isoforms in our new GBIG transcriptome (S4b).

### Differential expression during compensatory plasticity

Pairwise comparisons of normalized counts data from deafferented *vs*. control crickets were performed at each time point using the R package DESeq2 (See Supplemental Materials S5) using a False Discovery Rate (FDR) threshold of < 0.1 and an absolute value threshold for log2 fold changes of greater than 0.6. The distribution of differentially expressed genes was visualized using volcano plots (Figure 1 and with select points labeled in Supplemental Materials S6A-C). Of the few hundred genes identified as differentially expressed at 1-day post-deafferentation, only about 8% were downregulated 4-fold or more (14/173), while 29% were upregulated 4-fold or more (75/263). The remaining majority of significantly regulated genes at this early time point were up or downregulated more moderately at 2-4-fold (Figure 1A). Three days after deafferentation, 31% (22/71) were downregulated 4-fold or greater while 60% (131/218) were upregulated 4-fold or greater (Figure 1B). At seven days a relatively low number of genes were identified as differentially regulated, but the changes were large. Over 90% (13/14) were downregulated 4-fold or greater. More genes were upregulated rather than downregulated at seven days, but only 62.5% (15/24) of these were upregulated 4-fold or greater (Figure 1C).

**Figure 1.**
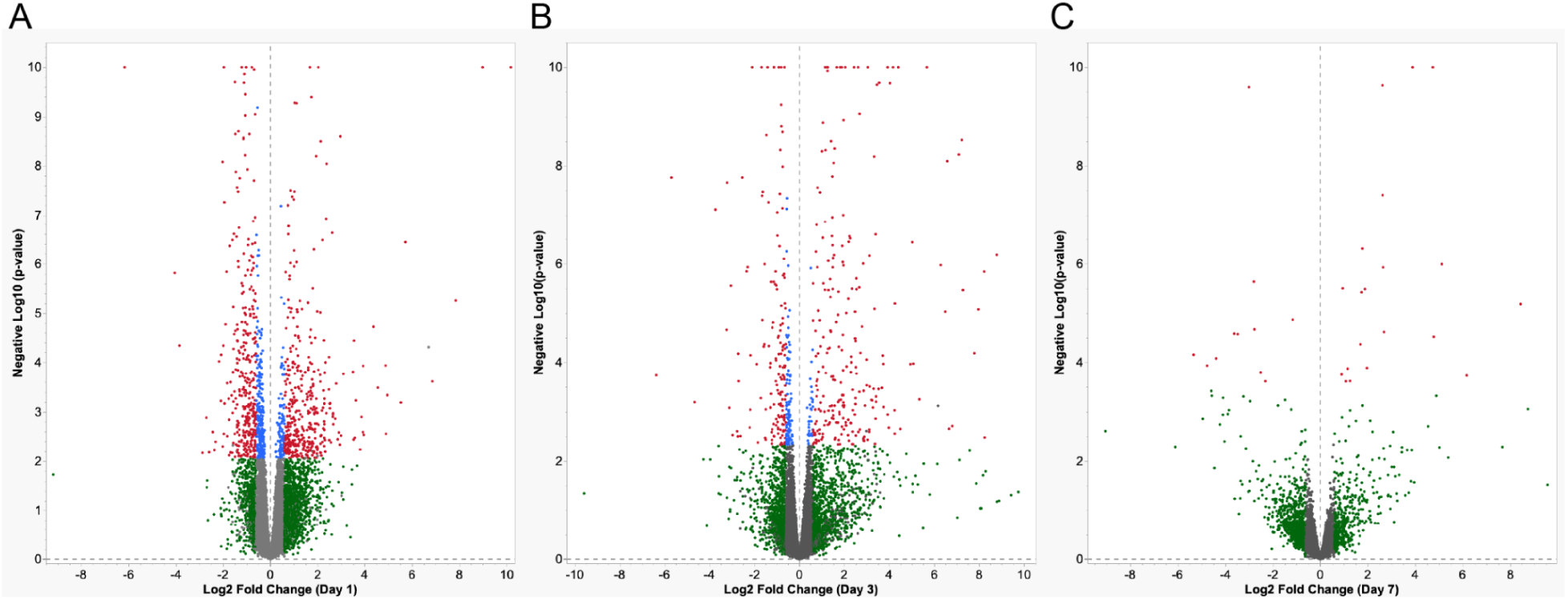
Volcano plots of differential gene expression in *G. bimaculatus* prothoracic ganglia at (A) one day, (B) 3 days, and (C) 7 days after deafferentation. Red dots represent genes that were determined to be differentially regulated by DESeq2, based on an absolute value of log2 fold change greater than 0.6 and an adjusted p-value less than 0.1. For visualization, all p-values less than 10^-10^ were set to 10^-10^. Blue dots show genes that were above threshold for adjusted p-value, but not log2 fold change. Green dots indicate genes that were above threshold for log2 fold change, but not adjusted p-value. (See Supplemental Figure S6 for volcano plots with gene names and species for most similar BLAST hits, where identified, for these significant genes.)

We used Venn diagrams to explore how many genes were differentially regulated across multiple days (Figure 2). The largest set of genes was upregulated at one day post-deafferentation (263), and 13 genes were uniquely shared between day one and day three (Figure 2A). Just over half of these 13 genes were unknown or uncharacterized genes (7/13); the genes that did have BLAST hits in this group included Beta-glucuronidase (GBI_01953), Dihydrofolate reductase (GBI_02419), Angiotensin-converting enzyme (GBI_15792), the Toll receptor Tollo (GBI_15807), Farnesol dehydrogenase (GBI_19142), and Esterase SG1 (GBIG_014732). Of the 24 genes upregulated at seven days, one gene, Follistatin (GBIG_021917), was also upregulated on day one. Five genes were uniquely upregulated at day three and day seven, and four of these are known: Embryonic Polarity Protein Dorsal (GBI_10428), the Mitochondrial Intermembrane Space Import Assembly Protein 40 (GBIG_008014), and two different Hexamerins (GBI_14215, GBI_14213). Three genes were upregulated across all three time points. One of these genes was identified as Uridine nucleosidase (GBI_16625), but the other two (GBI_16644, and GBI_161290) encode uncharacterized proteins (Figure 2A). When we examined the downregulated genes, we found the most to be downregulated on day 1 (173 genes) and that none were shared across multiple days (Figure 2B).

**Figure 2.**
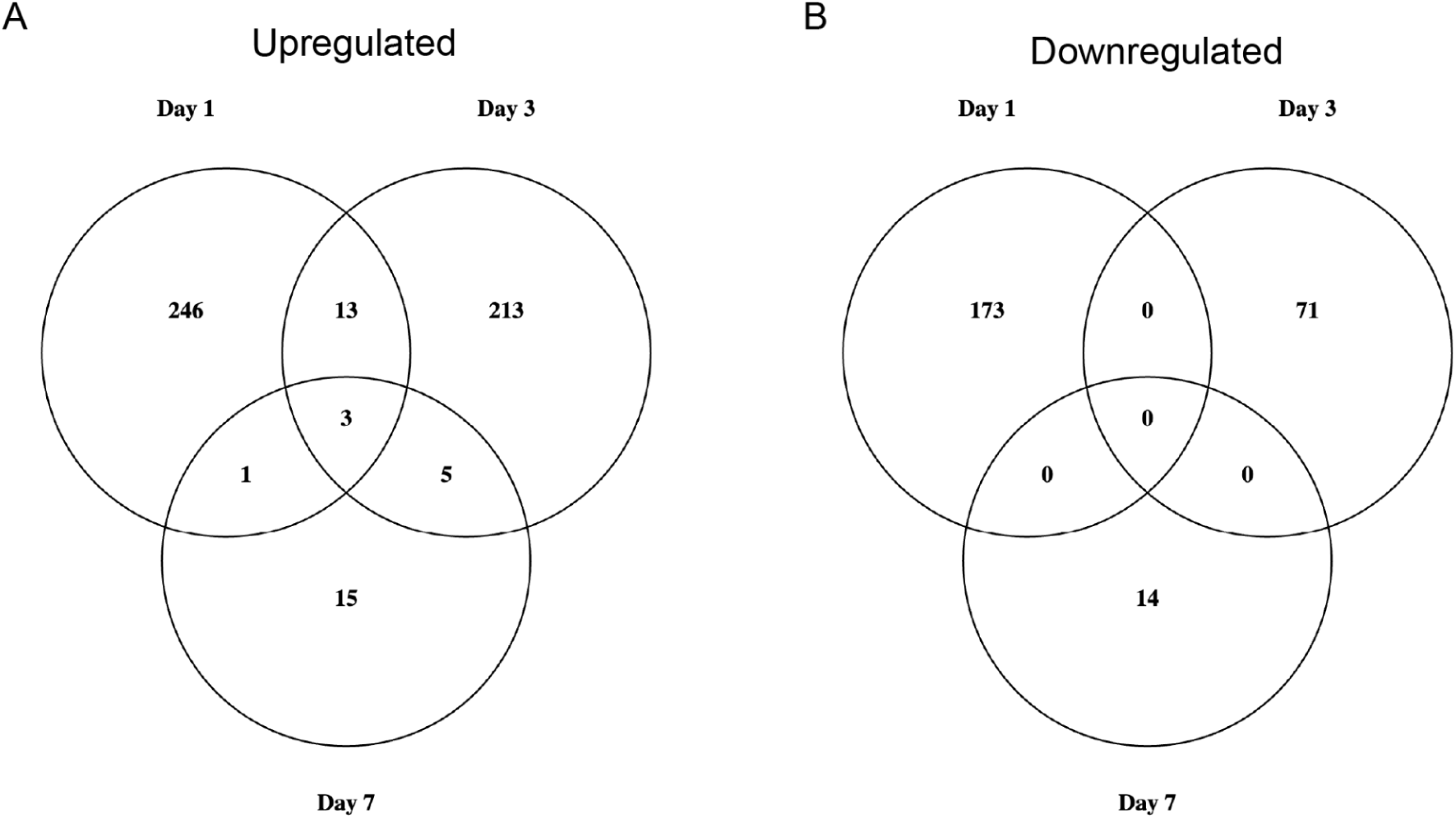
Venn diagrams show the total number of (A) upregulated genes and (B) downregulated genes across and between the three time points–one, three and seven days post-deafferentation. A small subset of genes were upregulated at multiple timepoints whereas downregulated genes showed no overlap across timepoints.

The ten transcripts with the largest fold-changes at each time point revealed that a majority (57%) were unidentified (Table 4). Of these 34 unidentified genes, 32% were uncharacterized or had no known function (orange), 22% showed no significant similarity to anything in the NCBI database (gray; determined by a 1e-10 threshold in BLAST-x), and 2 genes appear to have no open reading frames (“NA”).

**Table 4:** Top 10 upregulated and downregulated candidates by day after deafferentation. After further filtering out lowly expressed genes (<70), the highest-fold change genes are reported along with predicted protein length, log fold-change, adjusted p-value and BLAST-x results. The BLAST E-value threshold for proteins deemed to have no significant similarity (NSS) was >1e-10.

| Genome Assembly ID (>70) | Predicted Protein Length | Log Fold-Change | Adjusted p-Value | BLAST-x Result |
| --- | --- | --- | --- | --- |
| Day 1 Upregulated |  |  |  |  |
| GBI_04026 | 123 | 10.17769155 | 3.47E-12 | NSS |
| GBI_19006 | 2,127 | 6.854424676 | 0.00980655 | Vitellogenin-1 |
| GBI_00345 | 233 | 5.711323359 | 7.72857E-05 | Protein of Unknown Function |
| GBIG_021799 | 120 | 5.522997812 | 0.020181776 | NSS |
| GBIG_012536 | None | 4.947705344 | 0.01581712 | NA |
| GBI_09392 | 68 | 4.891409671 | 0.050533477 | Protein of Unknown Function |
| GBI_21898 | 169 | 4.549191474 | 0.012266513 | Myosin heavy chain |
| GBIG_008097 | 66 | 4.361993157 | 0.00168725 | NSS |
| GBI_22177 | 246 | 3.954731103 | 0.051945193 | Actin-87E |
| GBI_09921 | 526 | 3.923890575 | 0.006087742 | Maltase |
| Day 1 Downregulated |  |  |  |  |
| GBI_17240 | 707 | -6.17869514 | 5.25E-14 | Uncharacterized Protein |
| GBI_14155 | 185 | -4.049566288 | 0.00022171 | Protein of Unknown Function |
| GBI_03017 | 761 | -2.881235609 | 0.08408005 | Toll-like receptor Tollo |
| GBIG_020703 | 61 | -2.718103775 | 0.08408005 | NSS |
| GBI_11409 | 394 | -2.437291145 | 0.04822576 | Ionotropic receptor 883 |
| GBI_13795 | 1,747 | -2.323917701 | 0.06311496 | Chitinase 10 |
| GBI_16320 | 543 | -2.172279028 | 0.00541163 | Protein Wntless |
| GBI_07571 | 698 | -2.084224116 | 0.00434769 | Menin |
| GBI_09164 | 163 | -2.031558876 | 3.46E-06 | Protein of Unknown Function |
| GBIG_014182 | 121 | -2.025830779 | 0.07130012 | NSS |
| Day 3 Upregulated |  |  |  |  |
| GBIG_021611 | 304 | 9.337859769 | 7.63E-06 | Elongation of very long chain fatty acids protein |
| GBI_14312 | 577 | 8.224347814 | 0.00017188 | Uncharacterized Protein |
| GBI_09392 | 68 | 7.270907585 | 0.00034012 | Protein of Unknown Function |
| GBI_21041 | 74 | 7.229125563 | 9.28E-07 | Protein of Unknown Function |
| GBIG_013922 | 30 | 7.090303318 | 1.61E-06 | NSS |
| GBIG_021799 | 120 | 6.57102635 | 2.12E-06 | NSS |
| GBI_10707 | 334 | 6.483299448 | 0.00082479 | Regucalcin |
| GBIG_006506 | 513 | 5.664177412 | 7.82E-09 | Uncharacterized Protein |
| GBIG_000859 | 27 | 5.072274877 | 0.00626584 | Cubilin |
| GBI_04847 | 2,965 | 5.023596841 | 5.67E-05 | Cubilin |
| Day 3 Downregulated |  |  |  |  |
| GBIG_014798 | 69 | -6.376521814 | 0.00944352 | Uncharacterized Protein |
| GBI_16413 | 57 | -4.671154627 | 0.02511282 | NSS |
| GBIG_024533 | 500 | -3.734210908 | 1.4458E-05 | Uncharacterized Protein |
| GBI_13799 | 66 | -3.24358397 | 0.00171099 | Protein of Unknown Function |
| GBIG_002683 | 73 | -3.222328411 | 5.05E-06 | NSS |
| GBI_14967 | 57 | -3.116888272 | 0.029618 | Protein of Unknown Function |
| GBI_03346 | 814 | -3.052278026 | 0.00029659 | Serine protease grass |
| GBIG_001680 | 66 | -2.628150588 | 0.07650988 | NSS |
| GBI_11814 | 220 | -2.542921055 | 4.01E-06 | Protein obstructor-E |
| GBIG_004384 | 96 | -2.353731464 | 0.00017188 | No Significant Similarity |
| Day 7 Upregulated |  |  |  |  |
| GBI_15110 | 466 | 6.165429474 | 0.06701663 | Protein of Unknown Function |
| GBI_08082 | 541 | 5.115682556 | 0.00132352 | Cytochrome P450 4g15 |
| GBIG_006506 | 513 | 4.736256463 | 3.22E-27 | Protein of Unknown Function |
| GBIG_003347 | 80 | 3.883220428 | 8.27E-09 | NSS |
| GBIG_006623 | 701 | 3.845319594 | 2.39E-09 | Uncharacterized Protein |
| GBI_14213 | 205 | 3.56229324 | 0.0059807 | Hexamerin-like protein |
| GBI_14215 | 264 | 3.387846215 | 0.00902669 | Hexamerin |
| GBI_00444 | 229 | 2.644555038 | 0.00141071 | Hemolymph juvenile hormone-binding protein |
| GBI_21440 | 266 | 2.630198486 | 0.00418812 | Hexamerin-like protein |
| GBI_16129 | 197 | 2.622092223 | 6.4244E-05 | Protein of Unknown Function |
| Day 7 Downregulated |  |  |  |  |
| GBIG_005231 | None | -5.334741618 | 0.03254291 | NA |
| GBIG_023775 | 87 | 4.768617669 | 0.05085386 | NSS |
| GBIG_021273 | 959 | -4.383842295 | 0.0374451 | Retrovirus-related Pol polyprotein |
| GBI_10362 | 127 | -3.622861359 | 0.01458753 | Protein of Unknown Function |
| GBI_12769 | 436 | -3.484565574 | 0.01458753 | Cytochrome P450 |
| GBIG_024794 | 354 | -3.050121619 | 0.01960305 | Alpha amylase |
| GBIG_006668 | 65 | -3.00472941 | 4.81E-07 | NSS |
| GBI_11328 | 728 | -2.801729876 | 0.00249871 | Extracellular sulfatase |
| GBI_02488 | 565 | -2.510055292 | 0.0635878 | Alpha-amylase 2 |
| GBI_11696 | 784 | -2.212793974 | 0.08093477 | Uncharacterized Protein |

Of the genes with large-fold changes that could be identified, a few candidates were particularly notable. First, it was surprising to find such strong differential expression of several genes in this neuronal transcriptome that have not been previously associated with neurons, such as Vitellogenin (GBI_19006), Chitinase (GBI_13795), and Myosin heavy chain (GBI_21898). Vitellogenin is a lipid transport protein that functions as an egg yolk precursor protein, but is known to be expressed in glia in the central nervous system of honey bees (Münch et al., 2015) and likely regulates caste differentiation in those insects (Zhang et al., 2022). Chitinases may play a role in the support of the air-filled trachea (Merzendorfer and Zimoch, 2003), which branch through neuronal tissues. Chitinases also appear to have evolved a role in neuroinflammation in mammals and are currently being used as biomarkers for neurological disorders (Pinteac et al., 2021). Myosin heavy chain protein is a muscle-related gene identified in this analysis that was upregulated more than 20-fold, part of a larger group of differentially expressed muscle-related genes that we discuss below.

Several candidates in Table 2, including Regulcalcin (GBI_10707) and Alpha-amylase (GBI_02488), were identified by us in past suppression subtractive hybridization experiments (Horch et al., 2009). Though at the time we proposed a role in immune defense, stress response, and energy metabolism, we now know that Alpha amylase functions to degrade glycogen within synapses and is important for normal synaptic function (Byman et al., 2021). Regucalcin is important for calcium homeostasis and may protect against oxidative damage (Son et al., 2006). More recent results show that Regulcalcin may also provide resistance to oxidative stress, as has been specifically shown for amyloid-β toxicity in PC12 cells (Murata et al., 2018).

The greater than 4-fold downregulation of two additional genes, Wntless (GBI_16320) and Tollo (GBI_03017), at one day post-deafferentation was intriguing. The protein Wntless controls dendritic self-avoidance in *D melanogaster* and *C. elegans*. In the cricket auditory system, a rapid downregulation of Wntless could hypothetically alter the rules that typically guide dendrites and set the stage for the dendritic reorganization seen after deafferentation. Finally, GBI_03017, identified as Tollo (Toll-8), is likely a Toll receptor, but may be more similar to the clade of Toll 2’s rather than Toll 8 (data not shown.). Tolls are receptors for the family of Spaetzle ligands. Toll receptors are most commonly associated with immunity and dorsal-ventral patterning in early development (Lynch and Roth, 2011), but research in *D. melanogaster* suggests that Spz-Toll signaling may also have a neurotrophic-like role in neurons, regulating cell number, connectivity, and synaptogenesis (Anthoney et al., 2018).

### Protein-Protein Interaction Network Predictions

To gain a deeper understanding of the genes and predicted proteins identified in this analysis, we turned to STRING (Search Tool for the Retrieval of Interacting Genes/Proteins) analysis (Szklarczyk et al., 2023; see Methods), with a specific focus on GO Biological Process. We used the ranked-list analysis tool, sorting the DESeq2 output from each day on the basis of estimated log2 fold change, to identify functional groups of genes that were up- and/or down-regulated (Supplemental Material S7). The suggestion is that coordinated and biased expression of a functionally related set of genes indicates a systematic activation, deactivation, or regulation of the related processes. Visualizing the protein networks in our data that were systematically regulated can help suggest new hypotheses for the molecular basis of the anatomical plasticity seen in the cricket.

Based on this STRING analysis, we saw the enrichment of dozens of groups of functionally related genes on day one, mostly focused on signaling, membrane dynamics, and cellular migration (Figure 3). The enrichment of GTPase regulatory processes among downregulated genes was especially intriguing, because GTPAses are central to cytoskeletal rearrangements and because morphological changes in dendrites are evident only a few days post-deafferentation (Figure 3A). Though we had attempted to control for general injury-induced transcriptional changes by amputating the tarsus in our age-matched control animals (presumably a less-intrusive injury) many of the enriched proteins may be general responses to the injury

**Figure 3:**
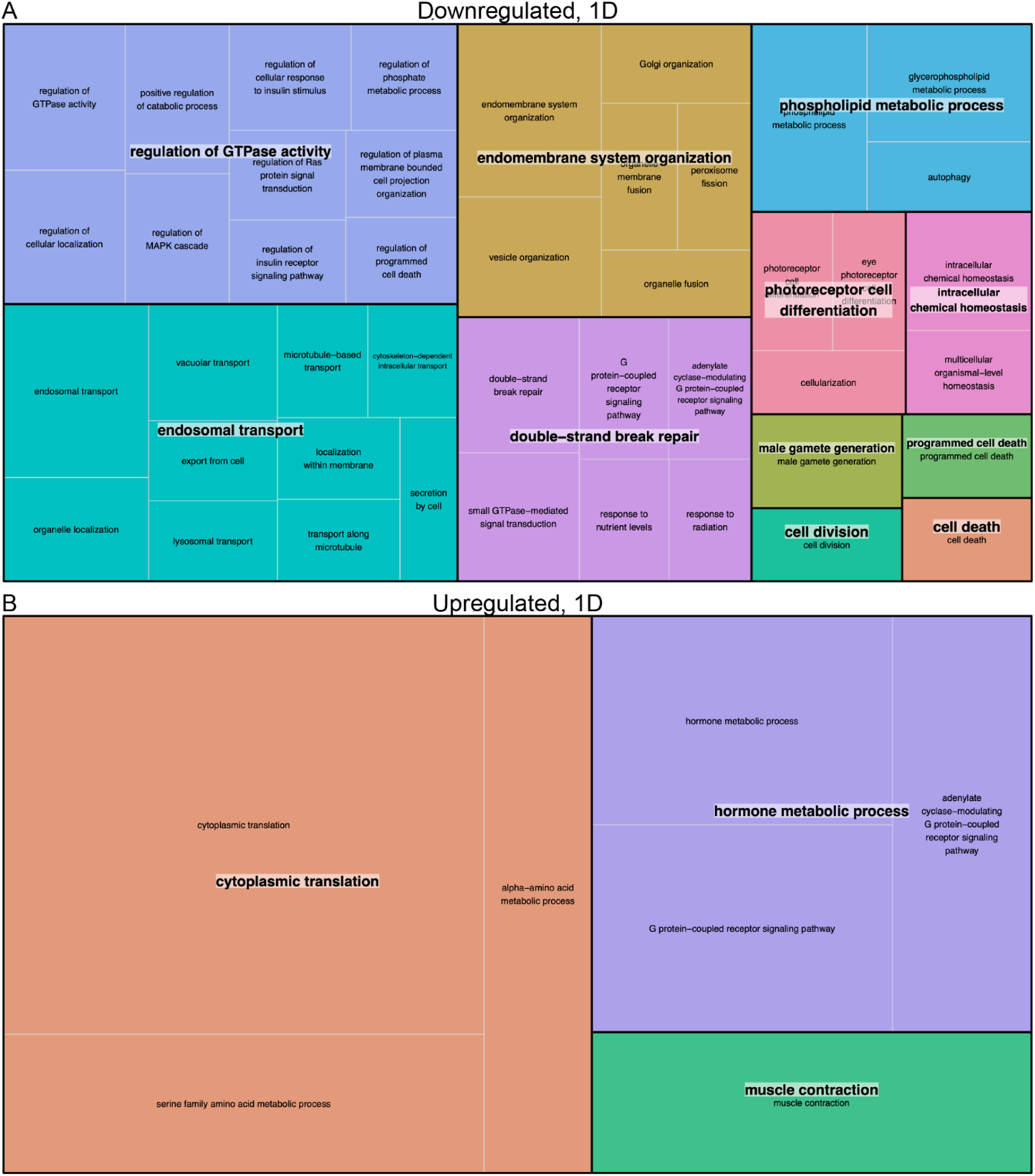
REVIGO TreeMap summary of gene ontology biological process terms identified as significantly (A) downregulated and (B) upregulated in the PTG one day after deafferentation. Each set of colored rectangles represents a group of semantically related terms with a common bold label. The size of the rectangles correlates with their relative p-values, with larger rectangles representing the GO terms with the highest level of significance and vice versa.

Given that we isolated exclusively prothoracic neuronal tissue for this transcriptome, one of the biggest surprises was the enrichment of muscle-related genes upregulated one day after deafferentation (Figure 3B), specifically in the clustered GO term “muscle contraction” (Figure 3B). Fifteen different transcripts, typically thought to be expressed predominantly or even exclusively in muscle, were enriched in our adult prothoracic ganglia after deafferentation, showing a PPI enrichment p-value of <1.0e-16 (Figure 4).

**Figure 4:**
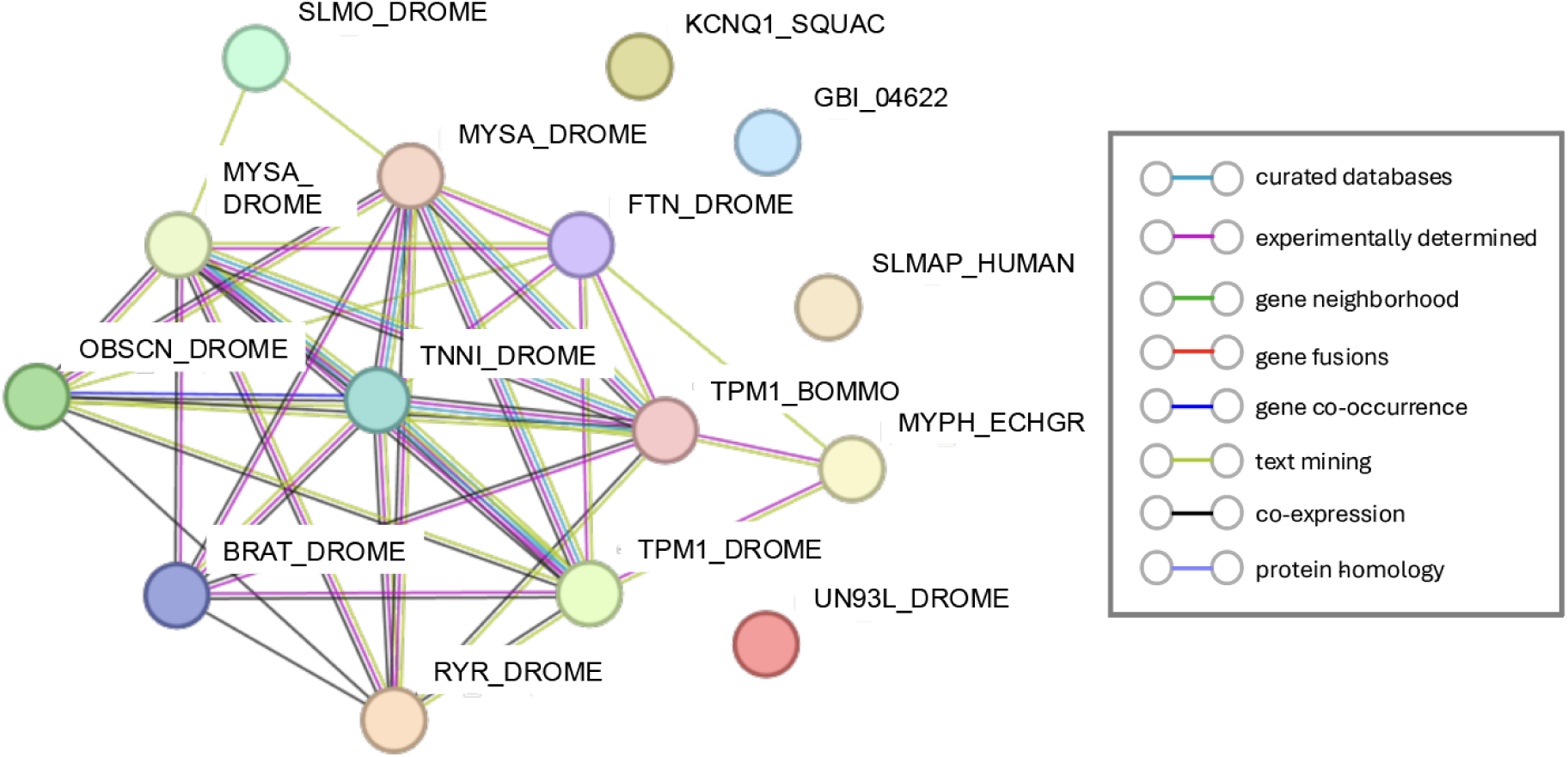
STRING diagram representing the protein interaction pathway derived from 15 muscle-related genes identified as enriched at one day post-deafferentation in our neuronal transcriptome. The average local clustering coefficient as reported by STRING was 0.486 and the protein-protein interaction (PPI) enrichment p-value was <1.0e-16. Known interactions from curated databases are shown in turquoise and interactional experimentally determined are shown in pink. Predicted interactions are shown in green (gene neighborhood), red (gene fusions), and purple (gene co-occurrence). Additional interactions are predicted from text mining (light green), co-expression (black), and protein homology (light purple).

Several of these proteins are thought to be exclusively expressed in muscle. For example Flightin (GBI_10829), first discovered in *D. melanogaster* flight muscles, was shown with immunoblots to be absent from other tissues (Vigoreaux et al., 1993). (We have successfully amplified *flightin* from cDNA created from PTG tissue, further confirming its presence in this neuronal tissue (data not shown)). On the other hand, many other proteins initially characterized in muscle have clearly been identified in neurons in other species, and several have important roles in growing neurons. For example, there is some limited evidence that Troponin (GBI_04010; Roisen et al., 1983), Tropomyosin (GBI_11216; Gray et al., 2017) and isoforms of Myosin heavy chain proteins (GBI_13123; GBI_21898; Rochlin et al., 1995) are expressed in neurons, and that they can function there to modulate neuronal morphology, including in the advancement and turning of growth cones (Gray et al., 2017; Rochlin et al., 1995). Our results raise the possibility that this set of proteins may have important roles to play in neurons, and possibly in the anatomical plasticity of neurons, beyond their known functions in muscle development and proliferation.

At three days post-deafferentation, based on the STRING analysis, it appears the deafferented neuronal tissue continues to adapt metabolically, while simultaneously engaging in stress response and cellular adaptation processes (Figure 5). Notably, hormone metabolic processes were enriched in both directions, indicating that these proteins might be at a high level of flux. The downregulation of proteins involved in synapse organization and transmission could be hallmarks of the ongoing dendritic reorganization that occurs around three days post-deafferentation. We know from time course studies that lateral AN dendrites are shortened while medial AN dendrites begin to sprout and extend across the midline, growing towards the contralateral axons (Horch et al., 2011).

**Figure 5:**
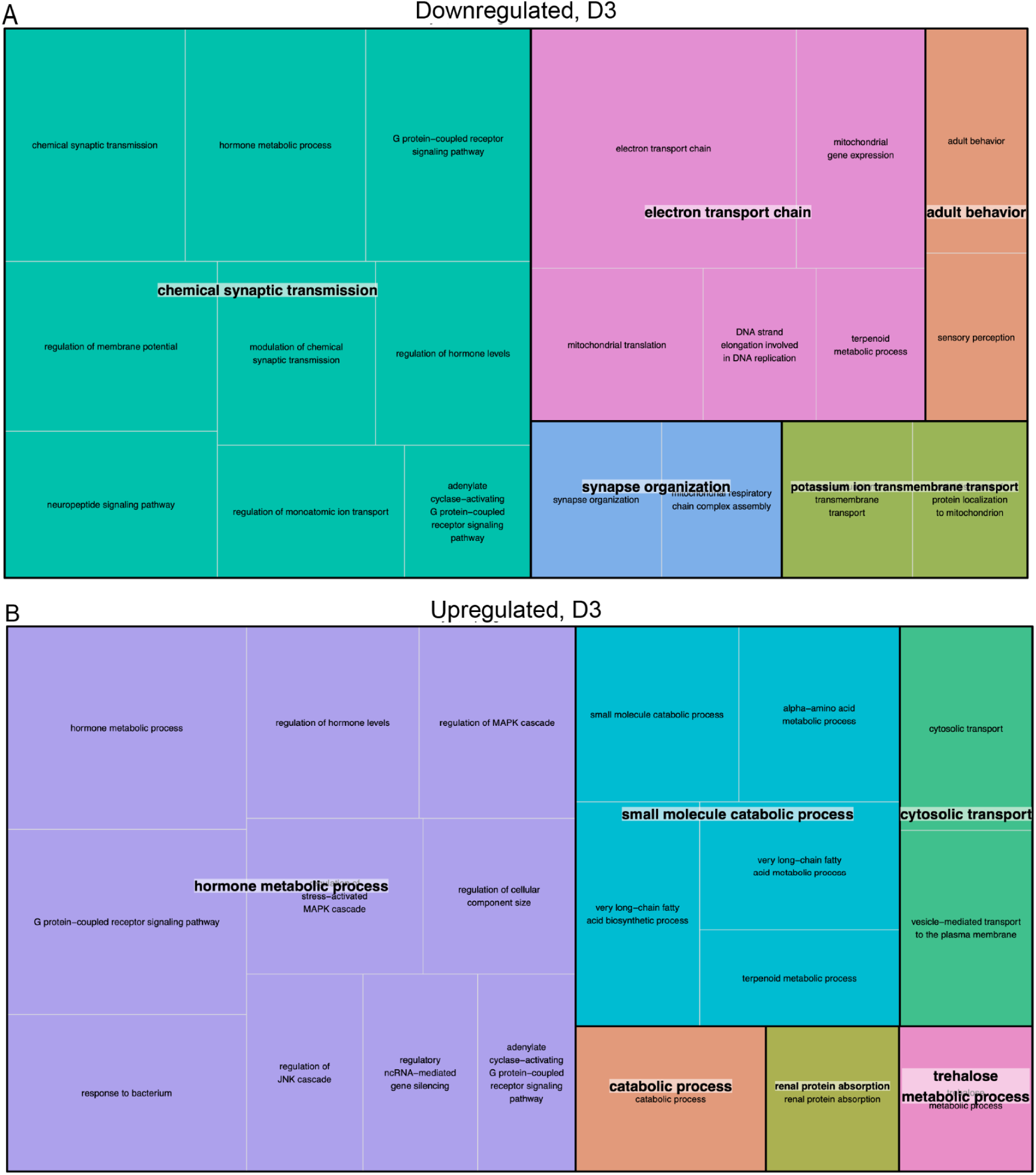
REVIGO treemap summary of gene ontology biological process terms that were significantly (A) downregulated and (B) upregulated in the PTG three days after deafferentation. Each set of colored rectangles represents a group of semantically related terms. The size of the rectangles correlates with their relative p-values, with larger rectangles representing the GO terms with the highest level of significance and vice versa.

Finally, at seven days, gave rise to only a few enriched GO terms as compared to days one and three. Most notable in this adult tissue, was the downregulation of genes related to the regulation of nervous system development. Of 193 known genes assigned to this process, 176 were mapped and were found to be skewed as a group towards downregulation with an FDR of 0.0029 (Supplemental Material S7). For example, this list included Semaphorins (Sema2a, Sema1a.1 and Sema1a.2) and their Plexin receptors, which we have characterized in the cricket (Horch et al., 2020; Maynard et al., 2007), several GTPases, as well as multiple factors that regulate GTPase function (Neurofibromin, Kalirin, Rheb, Rho-GAP-44, SAR1B, Tuberin). In addition, several proteins on this list influence Wnt signaling, which can influence synaptic development and plasticity (Budnik and Salinas, 2011). Proteins that influence synapse formation and axon guidance such as Enabled, FAK1, protein giant-lens, Inhibin, Kalirin, spartan, Tubulin polymerization-promoting protein, Calsyntenin-1 were also enriched. Also present were several factors that are known to influence dendritic formation and morphology, such as survival motor neuron protein1, the actin nucleation-promoting factor WASL, and the protocadherin-like wing polarity protein Flamingo/Starry Night (Gao et al., 2000).

### Conclusions

The injury-induced reorganization of auditory dendrites in the PTG of *Gryllus bimaculatus* is associated with the differential expression of hundreds of transcripts. This updated transcriptome, based on the integration of our RNAseq data with the reference genome annotation, added thousands of new genes and transcripts and therefore represents a more complete transcriptome than was previously published. Our findings highlight the importance of including a broad range of cellular tissue and phenotypic variation in the data used to define a transcriptome. We highlight transcriptional changes related to the regulation of GTPAse activity, muscle contraction, and nervous system development. The data presented here allows the development of targeted hypotheses that could elucidate the mechanisms responsible for the deafferentation-induced synaptic plasticity in the auditory system of crickets.

## Methods

### Animals, injury, and library preparation

Prothoracic ganglia from approximately 60 adult, male Mediterranean field crickets, *Gryllus bimaculatus,* were harvested and 21 individual ganglia were ultimately used as the sources of RNA for this transcriptome (Fisher et al., 2018). Male crickets that were adults for 3-5 days received either a control amputation of the distal segment of the left tarsus (“foot chop” control crickets), or the left prothoracic leg was severed mid-femur removing the auditory organ and deafferenting the ipsilateral central auditory neurons (“deafferented” experimental crickets). Males were chosen due to the potential sexual dimorphism in rates of dendritic growth after deafferentation (Pfister et al., 2013). Prothoracic ganglia were removed from crickets 1, 3, or 7 days after amputation at the femur or tarsus, or 18 hours post-backfill (Figure 6), and total RNA was purified as previously described (Fisher et al., 2018). In addition, several crickets were prepared for backfill as previously described (Horch et al. 2011). This tissue was sequenced for a different experiment, was used for the assembly but excluded from the differential expression analysis.

**Figure 6:**
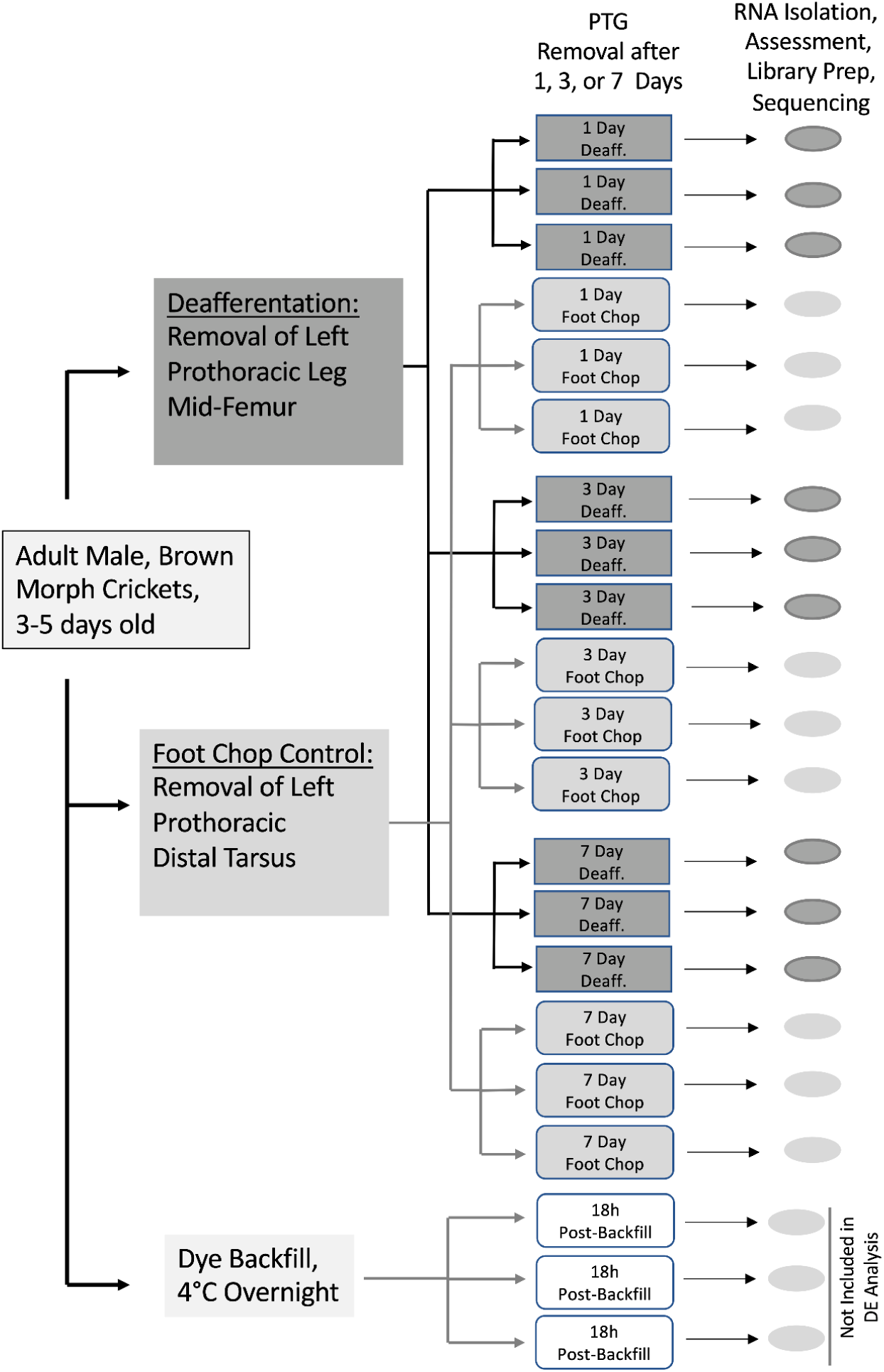
Summary of experimental design. 21 crickets, that were three to five days post adult eclosion, were amputated at the tarsal joint (“foot chop”) or mid-way along the femur (“deafferented”). PTGs were removed from deafferented or foot chop control animals one, three, or seven days post-injury. Three additional animals were backfilled 18 hrs prior to PTG removal; RNASeq data from these animals were included in the assembly but not in the differential expression.

The QIAGEN RNeasy Lipid Tissue Minikit was used to purify total RNA from each sample individually. RNA concentrations were assessed after TURBO DNA-free treatment (Ambion by Life Technologies) with a spectrophotometer (Nanodrop, Thermo Fisher Scientific) or a fluorometer (Qubit, Thermo Fisher Scientific). An Agilent 2100 Bioanalyzer (Applied Biosystems, Carlsbad, CA) was used to further assess sample quality. Based on evaluation of RNA quality and concentration of individual ganglion samples, the best 3 samples for each condition were selected for sequencing. Standard Illumina paired-end library protocols were used to prepare samples. The Illumina Hiseq 2500 platform, running v4 chemistry to generate ∼25M paired end reads of 100 bp in length for each sample, was used to sequence the RNA (Fisher et al., 2018).

### Transcriptome analysis and update

The draft cricket genome (*G. bimaculatus*) and the associated annotation GTF file were downloaded from https://gbimaculatusgenome.rc.fas.harvard.edu/ in June 2022. We used a newly developed Nextflow pipeline, txmupdate, that is designed to use empirical RNAseq data to update and revise a starting genome annotation [manuscript in preparation]. In brief, txmupdate was derived from the Nextflow (https://nextflow.io/) NF-core (https://nf-co.re/) rnaseq workflow, version 3.8.1 (https://nf-co.re/rnaseq/3.8.1), and the initial stages of quality control and alignment of individual samples to the genome with the STAR aligner are identical. Following alignment, each output BAM file was reduced using the program bamsifter, a utility that is part of the trinity rnaseq package (Haas et al., 2013), and then samtools merge (Li et al., 2009) was used to join the reduced files into a single unified BAM, which was finally again reduced with bamsifter. The final aggregate BAM (available here: https://doi.org/10.7910/DVN/EP0MXO) and with the associated .BAM.BAI file (S8)) was processed with stringtie (Pertea et al., 2015) to generate a sample-specific transcriptome, and the GFFcompare (Pertea and Pertea, 2020) was used to compare the sample-specific transcripts to the reference transcriptome. Finally, a novel program GTFinsert was used to join novel transcripts and the reference genome, resulting in a new updated GTF file (Supplemental File S1) describing the updated transcriptome.

### BUSCO transcriptome analysis

BUSCO (Benchmarking Universal Single Copy Orthologs) assessment (Manni et al., 2021a) of the transcriptome was performed using a Docker image of BUSCO version 5.7.0. Analyses were carried out against target set arthropoda_odb10 using the transcriptome analysis option. Final refinement was made by reconciling BUSCO matches from multiple transcript isoforms derived from a common gene resulting in the final “gene-level” tables presented here.

### Updated functional annotation of the GBIG transcriptome

In order to assess the likely functional roles of our updated transcriptome, we the following approach: First, the “annotate only” option for the TransPi workflow (Rivera-Vicéns et al., 2022), which calls the Trinotate pipeline, which is included in the TrinityRNAseq package (Grabherr et al., 2011). Trinotate assigns GO categories in three ways: (1) blastx (Camacho et al., 2009) alignments of the predicted transcripts, (2) blastp (Camacho et al., 2009) alignments of transdecoder-predicted proteins, and (3) pfam (Mistry et al., 2021) matches identified with hmmer (http://hmmer.org). Second, the transdecoder-predicted protein-coding sequences were reduced to the longest protein for each gene (Supplemental File S9). The complete set was uploaded to the STRING (Search Tool for the Retrieval of Interacting Genes/Proteins) database resource (Szklarczyk et al., 2023), which as of version 12 will process user submitted proteomes to provide the same annotations as are available to their natively hosted organisms. The TransPi analysis produced at least one GO annotation for 11,211 genes, the STRING analysis produced at least one GO annotation for 11,955 genes. All functional annotations for the updated genome are available as a Supplemental File (S10). The STRING-based tools for our updated transcriptome can be accessed at the STRING resource at URL https://version-12-0.string-db.org/organism/STRG0A58AIJ. Finally, any existing annotations in the reference GFF file were transferred directly to the new GTF.

### Differential gene expression analysis

Expression levels for each sample were generated for each original sample with the Nextflow Nf-core rnaseq pipeline (version 3.9) using the salmon pseudo alignment only option, with the updated transcriptome definition provided to define target transcripts.

Gene counts from Salmon were analyzed for differential gene expression using the R package “DESeq2” (version 1.42.1), performing a pairwise comparison between deafferented and control at each time point, generating estimated log2-fold change, p-value, and adjusted p-value, according to the Benjamini–Hochberg method (Benjamini and Hochberg, 1995). Genes were considered significantly up or down regulated if the False Discovery Rate (FDR) was greater than 0.1 and the absolute value of the log2 fold change was greater than 0.6. Differentially regulated gene lists for each day were used to make a Venn diagram in Venny 2.1 (https://bioinfogp.cnb.csic.es/tools/venny/)

### Functional enrichment analysis

To facilitate functional analysis, we used the “Annotated Proteome” feature, introduced in version 12 of the STRING database (Szklarczyk et al., 2025). The STRING analysis first searches the uploaded proteome against its existing database to identify putative orthologs. STRING then transfers protein-protein interaction links from other organisms, based on several different types of evidence to assess associations, combining the relevant information into a single confidence score for each association. We used TransDecoder [https://github.com/TransDecoder/TransDecoder] to generate a putative proteome file for our updated transcriptome, then reduced to a single protein per gene locus by arbitrarily selecting the longest protein predicted for each gene across all transcript isoforms. Functional enrichment was carried out through the STRING database interface (https://string-db.org/), using the uploaded cricket proteome as the target organism (https://version-12-0.string-db.org/organism/STRG0A58AIJ). Differential expression output from DESeq2 were threshold-selected to those with baseMean expression >= 100 counts and the resulting table of gene IDs and estimated log2 fold change were uploaded to STRING for use with the “Proteins with Ranks/Values” tool.

All searches of gene sets were carried out with default parameters. Gene Ontology Biological Process enrichment tables were downloaded from STRING and then segregated into up- and down-regulated terms, and subsequently submitted to the Revigo webserver interface (Supek et al., 2011) for generation of TreeMaps. The STRING ranked analysis is unusual in that, in addition to classifying groups of genes as skewed towards either up or downregulation, it will also identify groups that are skewed towards both ends. We added the categories classified as “both” into the up and downregulated categories before submission to Revigo. Final TreeMaps were generated by customization of the R script generated as part of the Revigo output.

## Supporting information

.BAM.BAI file (S8)

## Availability of data and material

Transcriptomic data are available on NCBI (Bioproject: PRJNA376023) (https://www.ncbi.nlm.nih.gov/bioproject/?term=PRJNA376023)

## Declarations

### Ethics approval and consent to participate

Not applicable.

### Authors’ contributions

HF, LL, and HWH conceived of and designed the experiments and HF collected and processed the tissue; FW, JHG, and JR completed the analysis; RAG and JHG performed the transcriptome update analysis; FW drafted the paper; HWH, and JHG substantially revised the manuscript. All authors read and approved the final manuscript

## Acknowledgements

We thank Marko Melendy for animal care support and Meera Prasad for consulting on revisions to this manuscript.

---

**S1 File: GTF file for all transcripts.**

**S2 File: FASTA file for all transcripts.**

**S3 Table: List of GBI genes joined together into a single GBIG gene.**

**S4 Figure: Examples of updated GBIG transcriptome annotations.** In each panel, the sashimi plot shows the count of reads supporting each splice-junction, based on the reduced BAM file generated with our data. The annotation plots below show GBI annotations (in black if present) and GBIG annotations (in blue). (A) The joining of two neighboring genes in the GBI annotations (GBI_10882 and GBI_10883) are supported by multiple spliced alignments that span portions of both genes. (B) Our transcriptome data and annotation process adds a critical new transcript to annotated gene GBI_00895. The novel transcript provides a BUSCO match that was not identified based on the GBI annotations. The inset shows an expanded view of four novel exons that allow for this annotation. (C) GBIG gene GBIG_020753, identified on genomic Scaffold56, is a novel identification with support for 11 distinct transcript isoforms.

**S5 File: DESeq2 output tables.**

**S6 Figure: Volcano plots with selected genes labeled for A) Day 1, B) Day 3, and C) Day 7.**

**S7 File: Biological process enrichment output from STRING for Day 1, 3, and 7.**

**S8 File: BAM.BAI file that is the index file for the .BAM data (**https://doi.org/10.7910/DVN/EP0MXO)

**S9 File: FASTA formatted file containing the longest predicted protein for each gene, which was uploaded to the STRING database.**

**S10 File: All annotations for the updated transcriptome assembly.**

## References

Anthoney, N., Foldi, I., Hidalgo, A., 2018. Toll and Toll-like receptor signalling in development. Development 145, dev156018. 10.1242/dev.156018

Bando, T., Ishimaru, Y., Kida, T., Hamada, Y., Matsuoka, Y., Nakamura, T., Ohuchi, H., Noji, S., Mito, T., 2013. Analysis of RNA-Seq data reveals involvement of JAK/STAT signalling during leg regeneration in the cricket Gryllus bimaculatus. Development 140, 959–964. 10.1242/dev.084590

Benjamini, Y., Hochberg, Y., 1995. Controlling the False Discovery Rate: A Practical and Powerful Approach to Multiple Testing. J. R. Stat. Soc. Ser. B Methodol. 57, 289–300.

Brodfuehrer, P.D., Hoy, R.R., 1988. Effect of auditory deafferentation on the synaptic connectivity of a pair of identified interneurons in adult field crickets. J. Neurobiol. 19, 17–38. 10.1002/neu.480190104

Budnik, V., Salinas, P.C., 2011. Wnt signaling during synaptic development and plasticity. Curr. Opin. Neurobiol., Developmental neuroscience 21, 151–159. 10.1016/j.conb.2010.12.002

Byman, E., Martinsson, I., Haukedal, H., Bank, T.N.B., Gouras, G., Freude, K.K., Wennström, M., 2021. Neuronal α-amylase is important for neuronal activity and glycogenolysis and reduces in presence of amyloid beta pathology. Aging Cell 20, e13433. 10.1111/acel.13433

Camacho, C., Coulouris, G., Avagyan, V., Ma, N., Papadopoulos, J., Bealer, K., Madden, T.L., 2009. BLAST+: architecture and applications. BMC Bioinformatics 10, 421. 10.1186/1471-2105-10-421

Fisher, H.P., Pascual, M.G., Jimenez, S.I., Michaelson, D.A., Joncas, C.T., Quenzer, E.D., Christie, A.E., Horch, H.W., 2018. De novo assembly of a transcriptome for the cricket Gryllus bimaculatus prothoracic ganglion: An invertebrate model for investigating adult central nervous system compensatory plasticity. PLOS ONE 13, e0199070. 10.1371/journal.pone.0199070

Gao, F.-B., Kohwi, M., Brenman, J.E., Jan, L.Y., Jan, Y.N., 2000. Control of Dendritic Field Formation in *Drosophila*. Neuron 28, 91–101. 10.1016/S0896-6273(00)00088-X

Grabherr, M.G., Haas, B.J., Yassour, M., Levin, J.Z., Thompson, D.A., Amit, I., Adiconis, X., Fan, L., Raychowdhury, R., Zeng, Q., Chen, Z., Mauceli, E., Hacohen, N., Gnirke, A., Rhind, N., di Palma, F., Birren, B.W., Nusbaum, C., Lindblad-Toh, K., Friedman, N., Regev, A., 2011. Trinity: reconstructing a full-length transcriptome without a genome from RNA-Seq data. Nat. Biotechnol. 29, 644–652. 10.1038/nbt.1883

Gray, K.T., Kostyukova, A.S., Fath, T., 2017. Actin regulation by tropomodulin and tropomyosin in neuronal morphogenesis and function. Mol. Cell. Neurosci. 84, 48–57. 10.1016/j.mcn.2017.04.002

Haas, B.J., Papanicolaou, A., Yassour, M., Grabherr, M., Blood, P.D., Bowden, J., Couger, M.B., Eccles, D., Li, B., Lieber, M., MacManes, M.D., Ott, M., Orvis, J., Pochet, N., Strozzi, F., Weeks, N., Westerman, R., William, T., Dewey, C.N., Henschel, R., LeDuc, R.D., Friedman, N., Regev, A., 2013. De novo transcript sequence reconstruction from RNA-seq using the Trinity platform for reference generation and analysis. Nat. Protoc. 8, 1494–1512. 10.1038/nprot.2013.084

Harel, N.Y., Strittmatter, S.M., 2006. Can regenerating axons recapitulate developmental guidance during recovery from spinal cord injury? Nat. Rev. Neurosci. 7, 603–616. 10.1038/nrn1957

Horch, H.W., McCarthy, S.S., Johansen, S.L., Harris, J.M., 2009. Differential gene expression during compensatory sprouting of dendrites in the auditory system of the cricket *Gryllus bimaculatus*. Insect Mol. Biol. 18, 483–496. 10.1111/j.1365-2583.2009.00891.x

Horch, H.W., Sheldon, E., Cutting, C.C., Williams, C.R., Riker, D.M., Peckler, H.R., Sangal, R.B., 2011. Bilateral Consequences of Chronic Unilateral Deafferentation in the Auditory System of the Cricket *Gryllus bimaculatus*. Dev. Neurosci. 33, 21–37. 10.1159/000322887

Horch, H.W., Spicer, S.B., Low, I.I.C., Joncas, C.T., Quenzer, E.D., Okoya, H., Ledwidge, L.M., Fisher, H.P., 2020. Characterization of *plexinA* and two distinct *semaphorin1a* transcripts in the developing and adult cricket *Gryllus bimaculatus*. J. Comp. Neurol. 528, 687–702. 10.1002/cne.24790

Hoy, R.R., Nolen, T.G., Casaday, G.C., 1985. Dendritic sprouting and compensatory synaptogenesis in an identified interneuron follow auditory deprivation in a cricket. Proc. Natl. Acad. Sci. 82, 7772–7776. 10.1073/pnas.82.22.7772

Kaneko, S., Iwanami, A., Nakamura, M., Kishino, A., Kikuchi, K., Shibata, S., Okano, H.J., Ikegami, T., Moriya, A., Konishi, O., Nakayama, C., Kumagai, K., Kimura, T., Sato, Y., Goshima, Y., Taniguchi, M., Ito, M., He, Z., Toyama, Y., Okano, H., 2006. A selective Sema3A inhibitor enhances regenerative responses and functional recovery of the injured spinal cord. Nat. Med. 12, 1380–1389. 10.1038/nm1505

Kerwin, S.K., Li, J.S.S., Noakes, P.G., Shin, G.J.-E., Millard, S.S., 2018. Regulated Alternative Splicing of Drosophila Dscam2 Is Necessary for Attaining the Appropriate Number of Photoreceptor Synapses. Genetics 208, 717–728. 10.1534/genetics.117.300432

Li, H., Handsaker, B., Wysoker, A., Fennell, T., Ruan, J., Homer, N., Marth, G., Abecasis, G., Durbin, R., 1000 Genome Project Data Processing Subgroup, 2009. The Sequence Alignment/Map format and SAMtools. Bioinformatics 25, 2078–2079. 10.1093/bioinformatics/btp352

Li, J.S.S., Millard, S.S., 2019. Deterministic splicing of Dscam2 is regulated by Muscleblind. Sci. Adv. 5, eaav1678. 10.1126/sciadv.aav1678

Lynch, J.A., Roth, S., 2011. The evolution of dorsal–ventral patterning mechanisms in insects. Genes Dev. 25, 107–118. 10.1101/gad.2010711

Manni, M., Berkeley, M.R., Seppey, M., Simão, F.A., Zdobnov, E.M., 2021a. BUSCO Update: Novel and Streamlined Workflows along with Broader and Deeper Phylogenetic Coverage for Scoring of Eukaryotic, Prokaryotic, and Viral Genomes. Mol. Biol. Evol. 38, 4647–4654. 10.1093/molbev/msab199

Manni, M., Berkeley, M.R., Seppey, M., Zdobnov, E.M., 2021b. BUSCO: Assessing Genomic Data Quality and Beyond. Curr. Protoc. 1, e323. 10.1002/cpz1.323

Maynard, K.R., McCarthy, S.S., Sheldon, E., Horch, H.W., 2007. Developmental and adult expression of semaphorin 2a in the cricket *Gryllus bimaculatus*. J. Comp. Neurol. 503, 169–181. 10.1002/cne.21392

Merzendorfer, H., Zimoch, L., 2003. Chitin metabolism in insects: structure, function and regulation of chitin synthases and chitinases. J. Exp. Biol. 206, 4393–4412. 10.1242/jeb.00709

Mistry, J., Chuguransky, S., Williams, L., Qureshi, M., Salazar, G.A., Sonnhammer, E.L.L., Tosatto, S.C.E., Paladin, L., Raj, S., Richardson, L.J., Finn, R.D., Bateman, A., 2021. Pfam: The protein families database in 2021. Nucleic Acids Res. 49, D412–D419. 10.1093/nar/gkaa913

Münch, D., Ihle, K.E., Salmela, H., Amdam, G.V., 2015. Vitellogenin in the honey bee brain: Atypical localization of a reproductive protein that promotes longevity. Exp. Gerontol., Aging in the Wild: Insights from Free-Living and Non-Model organisms 71, 103–108. 10.1016/j.exger.2015.08.001

Murata, T., Yamaguchi, M., Kohno, S., Takahashi, C., Kakimoto, M., Sugimura, Y., Kamihara, M., Hikita, K., Kaneda, N., 2018. Regucalcin confers resistance to amyloid-β toxicity in neuronally differentiated PC 12 cells. FEBS Open Bio 8, 349–360. 10.1002/2211-5463.12374

Pertea, G., Pertea, M., 2020. GFF Utilities: GffRead and GffCompare. F1000Research 9, ISCB Comm J-304. 10.12688/f1000research.23297.2

Pertea, M., Pertea, G.M., Antonescu, C.M., Chang, T.-C., Mendell, J.T., Salzberg, S.L., 2015. StringTie enables improved reconstruction of a transcriptome from RNA-seq reads. Nat. Biotechnol. 33, 290–295. 10.1038/nbt.3122

Pfister, A., Johnson, A., Ellers, O., Horch, H.W., 2013. Quantification of dendritic and axonal growth after injury to the auditory system of the adult cricket Gryllus bimaculatus. Front. Physiol. 3. 10.3389/fphys.2012.00367

Pinteac, R., Montalban, X., Comabella, M., 2021. Chitinases and chitinase-like proteins as biomarkers in neurologic disorders. Neurol. Neuroimmunol. Neuroinflammation 8, e921. 10.1212/NXI.0000000000000921

Popov, A.V., Markovich, A.M., Andjan, A.S., 1978. Auditory interneurons in the prothoracic ganglion of the cricket, gryllus bimaculatus deGeer: I. The large segmental auditory neuron (LSAN). J. Comp. Physiol. A 126, 183–192. 10.1007/BF00666372

Poulet, J.F.A., Hedwig, B., 2001. Tympanic membrane oscillations and auditory receptor activity in the stridulating cricket Gryllus bimaculatus. J. Exp. Biol. 204, 1281–1293. 10.1242/jeb.204.7.1281

Prasad, M.P., Detchou, D.K.E., Wang, F., Ledwidge, L.L., Kingston, S.E., Wilson Horch, H., 2021. Transcriptional expression changes during compensatory plasticity in the terminal ganglion of the adult cricket Gryllus bimaculatus. BMC Genomics 22, 742. 10.1186/s12864-021-08018-x

Prigge, C.L., Kay, J.N., 2018. Dendrite morphogenesis from birth to adulthood. Curr. Opin. Neurobiol. 53, 139–145. 10.1016/j.conb.2018.07.007

Rivera-Vicéns, R.E., Garcia-Escudero, C.A., Conci, N., Eitel, M., Wörheide, G., 2022. TransPi—a comprehensive TRanscriptome ANalysiS PIpeline for de novo transcriptome assembly. Mol. Ecol. Resour. 22, 2070–2086. 10.1111/1755-0998.13593

Rivera-Vicéns, R.E., Garcia-Escudero, C.A., Conci, N., Eitel, M., Wörheide, G., 2022. TransPi—a comprehensive TRanscriptome ANalysiS PIpeline for *de novo* transcriptome assembly. Mol. Ecol. Resour. 22, 2070–2086. 10.1111/1755-0998.13593

Rochlin, M.W., Itoh, K., Adelstein, R.S., Bridgman, P.C., 1995. Localization of myosin II A and B isoforms in cultured neurons. J. Cell Sci. 108, 3661–3670. 10.1242/jcs.108.12.3661

Roisen, F.J., Wilson, F.J., Yorke, G., Inczedy-Marcsek, M., Hirabayashi, T., 1983. Immunohistochemical localization of troponin-C in cultured neurons. J. Muscle Res. Cell Motil. 4, 163–175. 10.1007/BF00712028

Sampaio-Baptista, C., Sanders, Z.-B., Johansen-Berg, H., 2018. Structural Plasticity in Adulthood with Motor Learning and Stroke Rehabilitation. Annu. Rev. Neurosci. 41, 25–40. 10.1146/annurev-neuro-080317-062015

Schildberger, K., Wohlers, D.W., Schmitz, B., Kleindienst, H.-U., Huber, F., 1986. Morphological and physiological changes in central auditory neurons following unilateral foreleg amputation in larval crickets. J. Comp. Physiol. A 158, 291–300. 10.1007/BF00603613

Schmitz, B., 1989. Neuroplasticity and phonotaxis in monaural adult female crickets (Gryllus bimaculatus de Geer). J. Comp. Physiol. A 164, 343–358. 10.1007/BF00612994

Schmucker, D., Clemens, J.C., Shu, H., Worby, C.A., Xiao, J., Muda, M., Dixon, J.E., Zipursky, S.L., 2000. Drosophila Dscam Is an Axon Guidance Receptor Exhibiting Extraordinary Molecular Diversity. Cell 101, 671–684. 10.1016/S0092-8674(00)80878-8

Simão, F.A., Waterhouse, R.M., Ioannidis, P., Kriventseva, E.V., Zdobnov, E.M., 2015. BUSCO: assessing genome assembly and annotation completeness with single-copy orthologs. Bioinformatics 31, 3210–3212. 10.1093/bioinformatics/btv351

Son, T.G., Zou, Y., Jung, K.J., Yu, B.P., Ishigami, A., Maruyama, N., Lee, J., 2006. SMP30 deficiency causes increased oxidative stress in brain. Mech. Ageing Dev. 127, 451–457. 10.1016/j.mad.2006.01.005

Supek, F., Bošnjak, M., Škunca, N., Šmuc, T., 2011. REVIGO summarizes and visualizes long lists of gene ontology terms. PloS One 6, e21800. 10.1371/journal.pone.0021800

Szklarczyk, D., Kirsch, R., Koutrouli, M., Nastou, K., Mehryary, F., Hachilif, R., Gable, A.L., Fang, T., Doncheva, N.T., Pyysalo, S., Bork, P., Jensen, L.J., von Mering, C., 2023. The STRING database in 2023: protein–protein association networks and functional enrichment analyses for any sequenced genome of interest. Nucleic Acids Res. 51, D638–D646. 10.1093/nar/gkac1000

Szklarczyk, D., Nastou, K., Koutrouli, M., Kirsch, R., Mehryary, F., Hachilif, R., Hu, D., Peluso, M.E., Huang, Q., Fang, T., Doncheva, N.T., Pyysalo, S., Bork, P., Jensen, L.J., von Mering, C., 2025. The STRING database in 2025: protein networks with directionality of regulation. Nucleic Acids Res. 53, D730–D737. 10.1093/nar/gkae1113

Vigoreaux, J.O., Saide, J.D., Valgeirsdottir, K., Pardue, M.L., 1993. Flightin, a novel myofibrillar protein of Drosophila stretch-activated muscles. J. Cell Biol. 121, 587–598. 10.1083/jcb.121.3.587

Wohlers, D.W., Huber, F., 1985. Topographical organization of the auditory pathway within the prothoracic ganglion of the cricket Gryllus campestris L. Cell Tissue Res. 239, 555–565. 10.1007/BF00219234

Ylla, G., Nakamura, T., Itoh, T., Kajitani, R., Toyoda, A., Tomonari, S., Bando, T., Ishimaru, Y., Watanabe, T., Fuketa, M., Matsuoka, Y., Barnett, A.A., Noji, S., Mito, T., Extavour, C.G., 2021a. Insights into the genomic evolution of insects from cricket genomes. Commun. Biol. 4, 1–12. 10.1038/s42003-021-02197-9

Ylla, G., Nakamura, T., Itoh, T., Kajitani, R., Toyoda, A., Tomonari, S., Bando, T., Ishimaru, Y., Watanabe, T., Fuketa, M., Matsuoka, Y., Barnett, A.A., Noji, S., Mito, T., Extavour, C.G., 2021b. Insights into the genomic evolution of insects from cricket genomes. Commun. Biol. 4, 1–12. 10.1038/s42003-021-02197-9

Yu, H.-H., Araj, H.H., Ralls, S.A., Kolodkin, A.L., 1998. The Transmembrane Semaphorin Sema I Is Required in Drosophila for Embryonic Motor and CNS Axon Guidance. Neuron 20, 207–220. 10.1016/S0896-6273(00)80450-X

Zeng, V., Ewen-Campen, B., Horch, H.W., Roth, S., Mito, T., Extavour, C.G., 2013. Developmental Gene Discovery in a Hemimetabolous Insect: De Novo Assembly and Annotation of a Transcriptome for the Cricket Gryllus bimaculatus. PLoS ONE 8, e61479. 10.1371/journal.pone.0061479

Zeng, V., Extavour, C.G., 2012. ASGARD: an open-access database of annotated transcriptomes for emerging model arthropod species. Database J. Biol. Databases Curation 2012, bas048. 10.1093/database/bas048

Zhang, W., Wang, L., Zhao, Y., Wang, Y., Chen, C., Hu, Y., Zhu, Y., Sun, H., Cheng, Y., Sun, Q., Zhang, J., Chen, D., 2022. Single-cell transcriptomic analysis of honeybee brains identifies vitellogenin as caste differentiation-related factor. iScience 25. 10.1016/j.isci.2022.104643

