## Supplemental Figure (S4) for "Integration of a neuronal RNAseq dataset with the draft *Gryllus bimaculatus* transcriptome refines gene predictions and highlights potential systematic response to injury"

### Slide 1
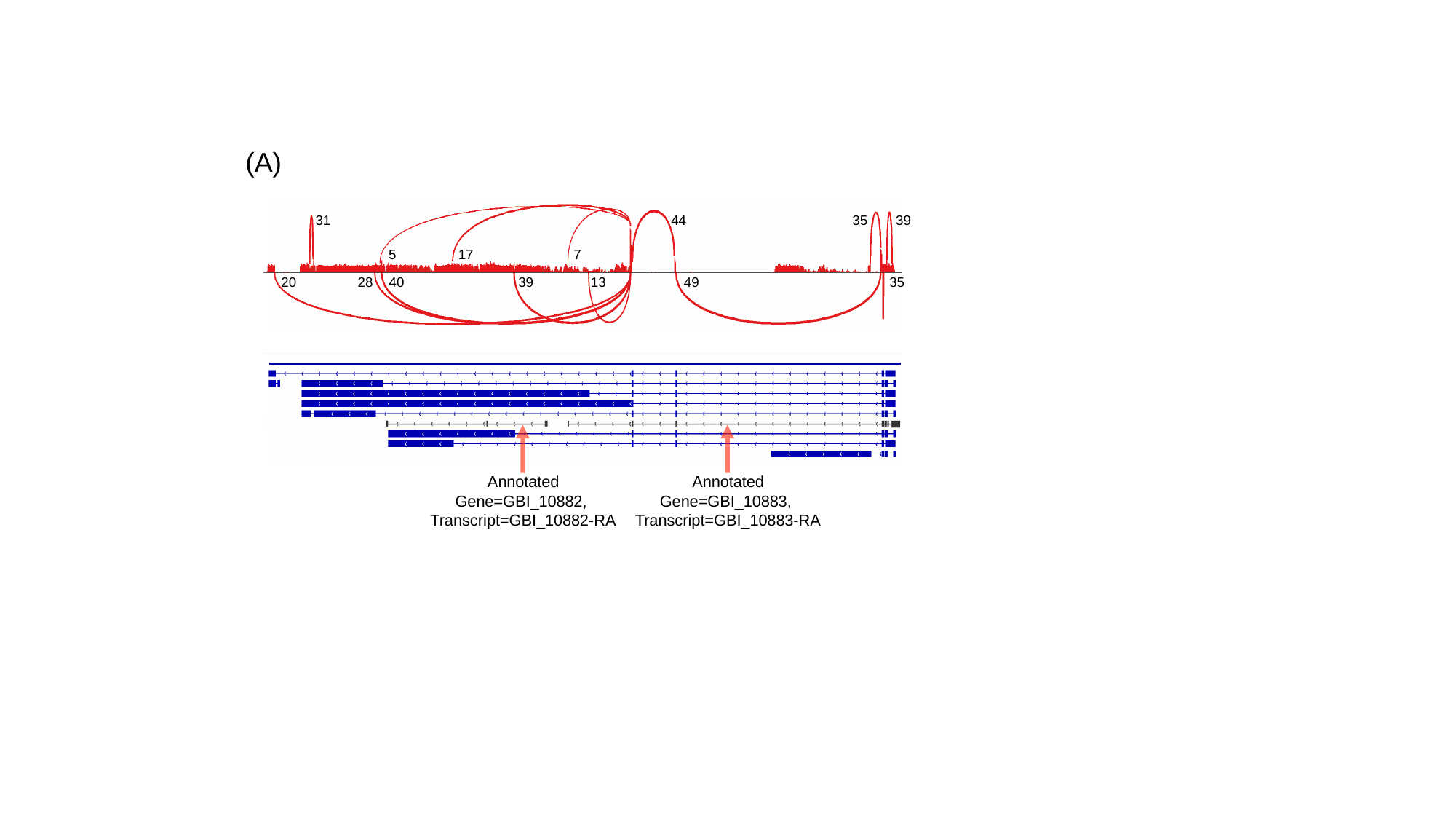

(A)
31
44
35
39
5
17
7
20
28
40
39
13
49
35
Annotated
Gene=GBI_10883,
Transcript=GBI_10883-RA
Annotated
Gene=GBI_10882,
Transcript=GBI_10882-RA

### Slide 2
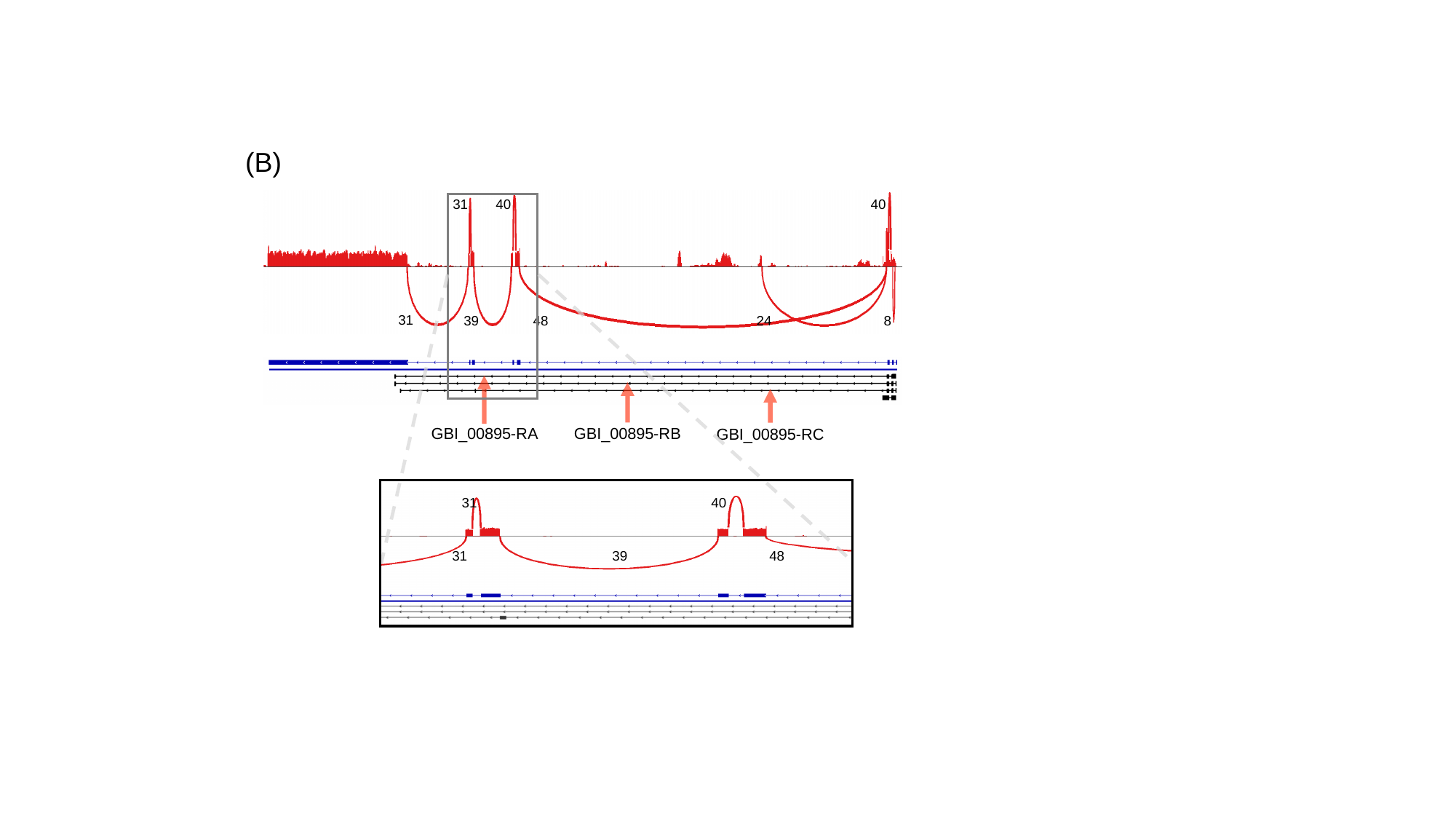

(B)
31
40
40
31
39
48
24
8
GBI_00895-RA
GBI_00895-RB
GBI_00895-RC
31
40
31
39
48

### Slide 3
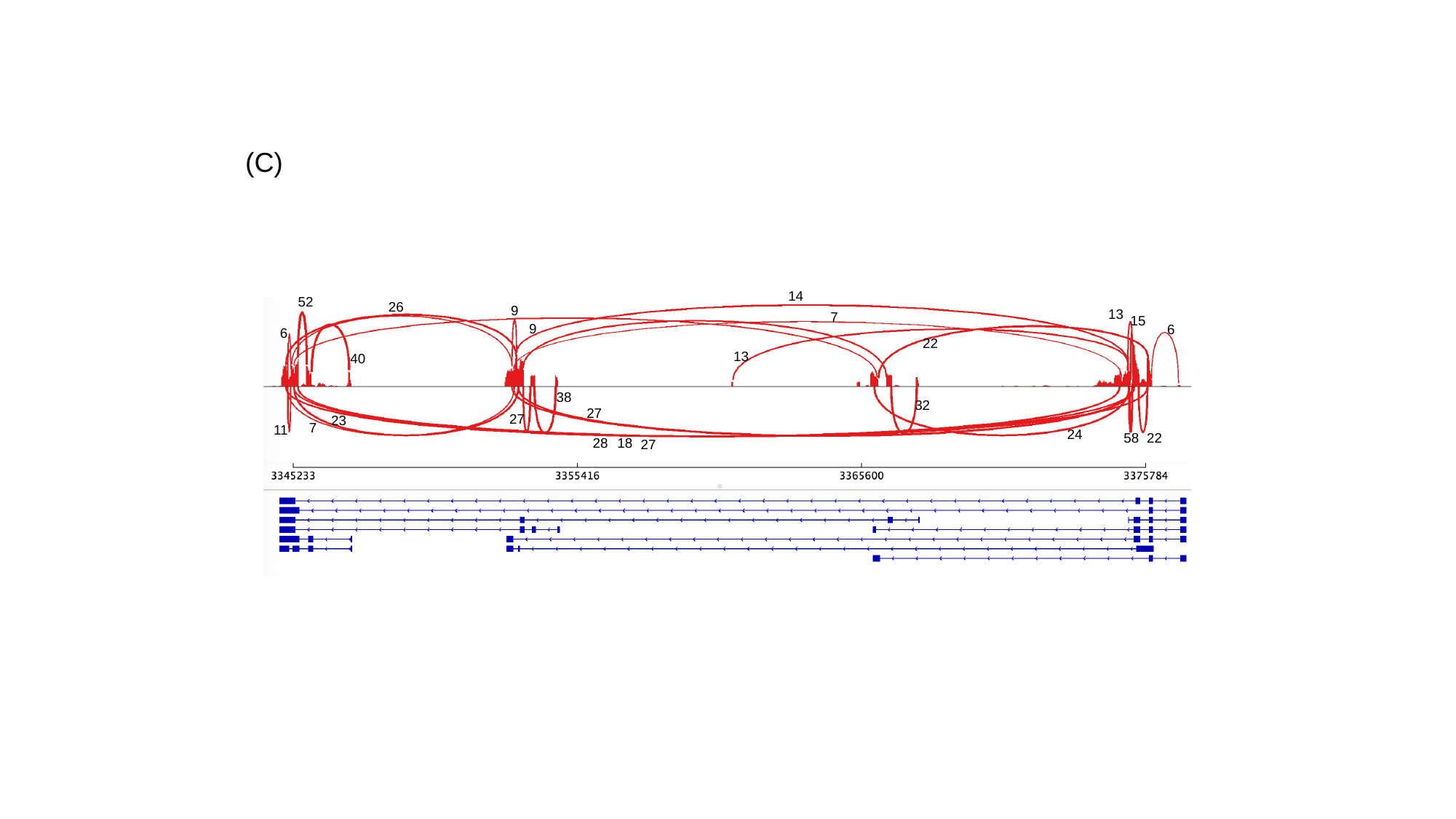

(C)
14
52
26
9
13
7
15
9
6
6
22
13
40
38
32
27
27
23
7
11
24
22
58
18
28
27

### Slide 4
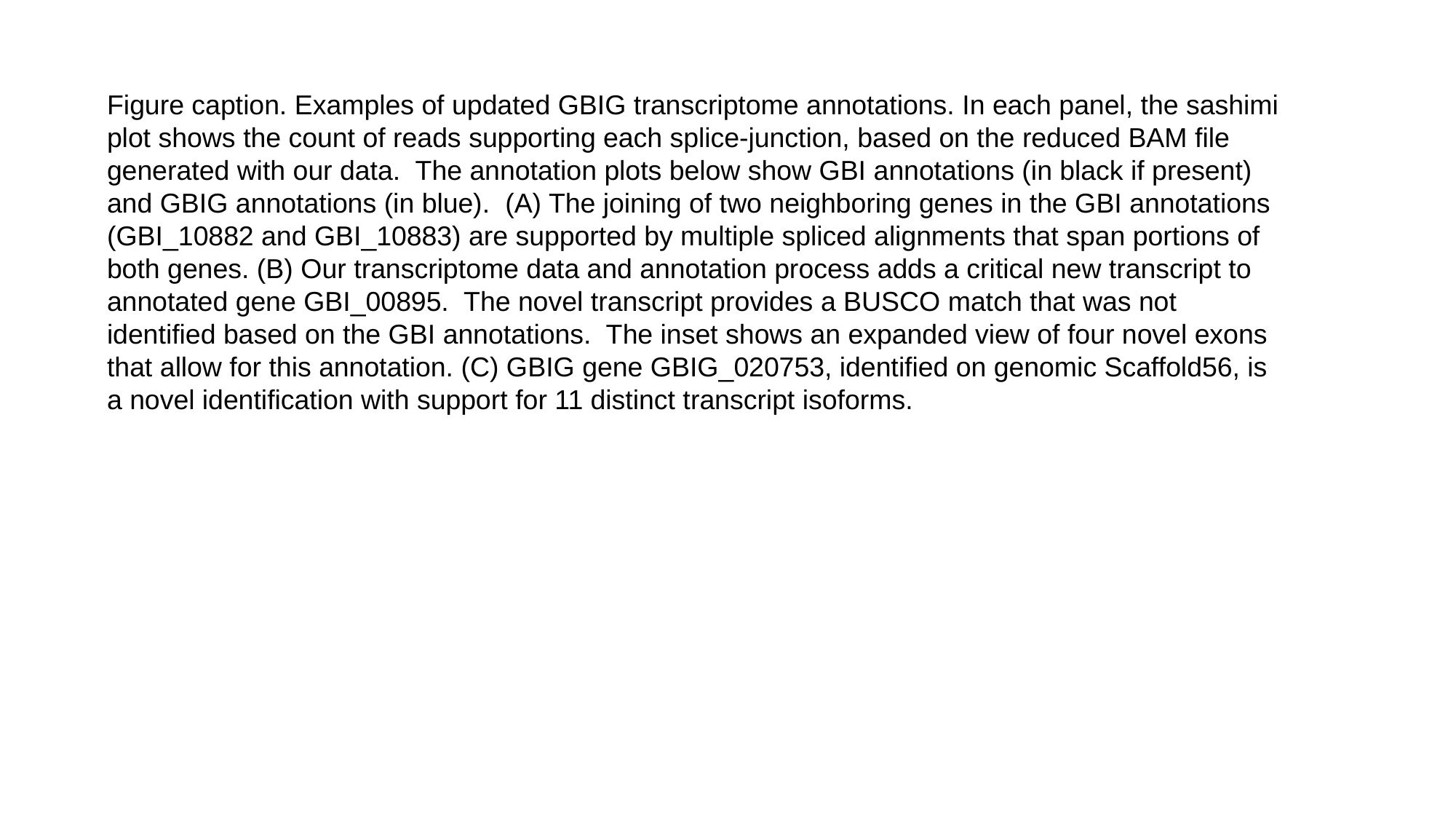

Figure caption. Examples of updated GBIG transcriptome annotations. In each panel, the sashimi plot shows the count of reads supporting each splice-junction, based on the reduced BAM file generated with our data. The annotation plots below show GBI annotations (in black if present) and GBIG annotations (in blue). (A) The joining of two neighboring genes in the GBI annotations (GBI_10882 and GBI_10883) are supported by multiple spliced alignments that span portions of both genes. (B) Our transcriptome data and annotation process adds a critical new transcript to annotated gene GBI_00895. The novel transcript provides a BUSCO match that was not identified based on the GBI annotations. The inset shows an expanded view of four novel exons that allow for this annotation. (C) GBIG gene GBIG_020753, identified on genomic Scaffold56, is a novel identification with support for 11 distinct transcript isoforms.
