## Supplementary figures and images for "Integration of a neuronal RNAseq dataset with the draft *Gryllus bimaculatus* transcriptome refines gene predictions and highlights potential systematic response to injury"

### Supplemental Materials S6

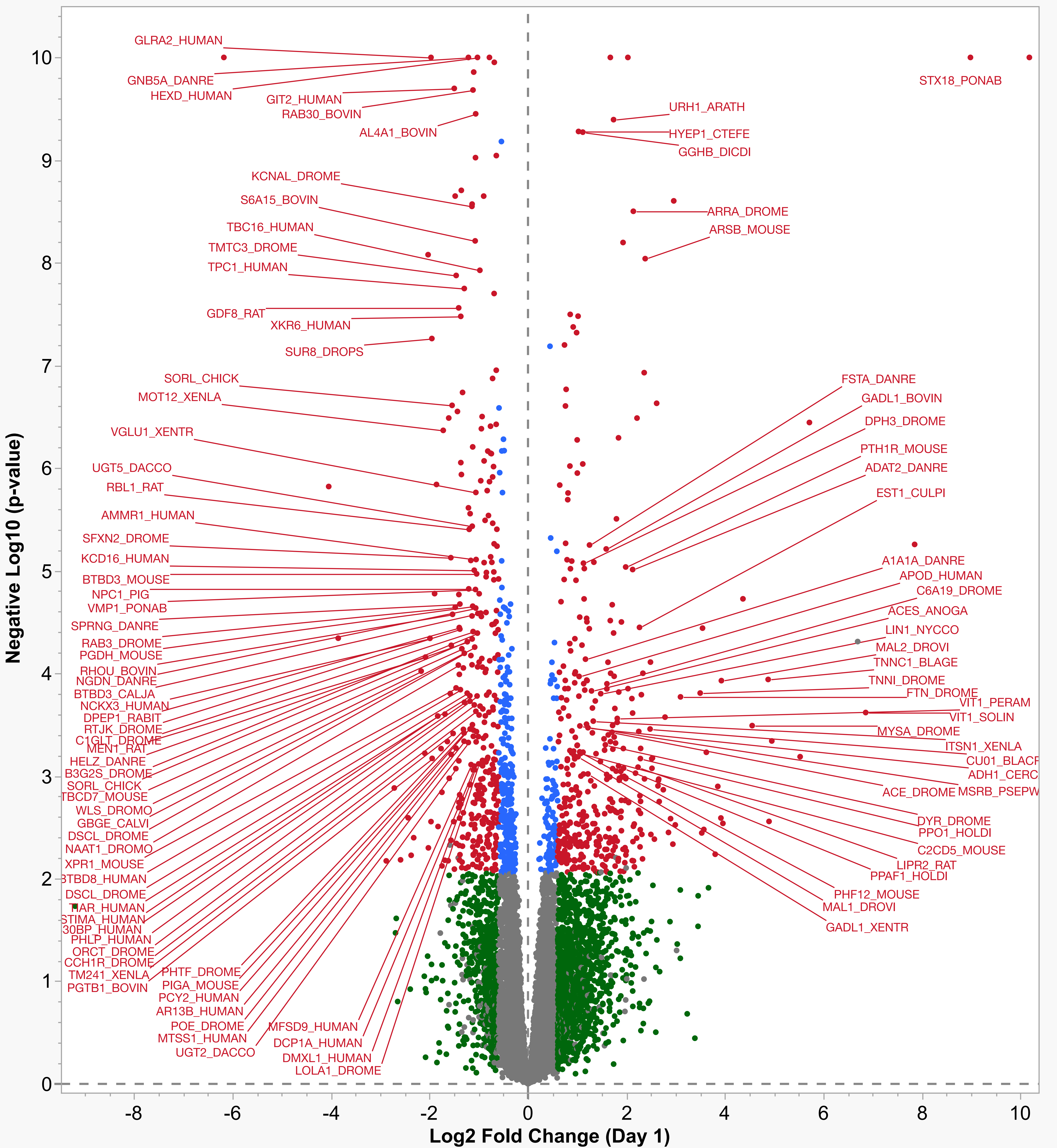

### Supplemental Materials S6

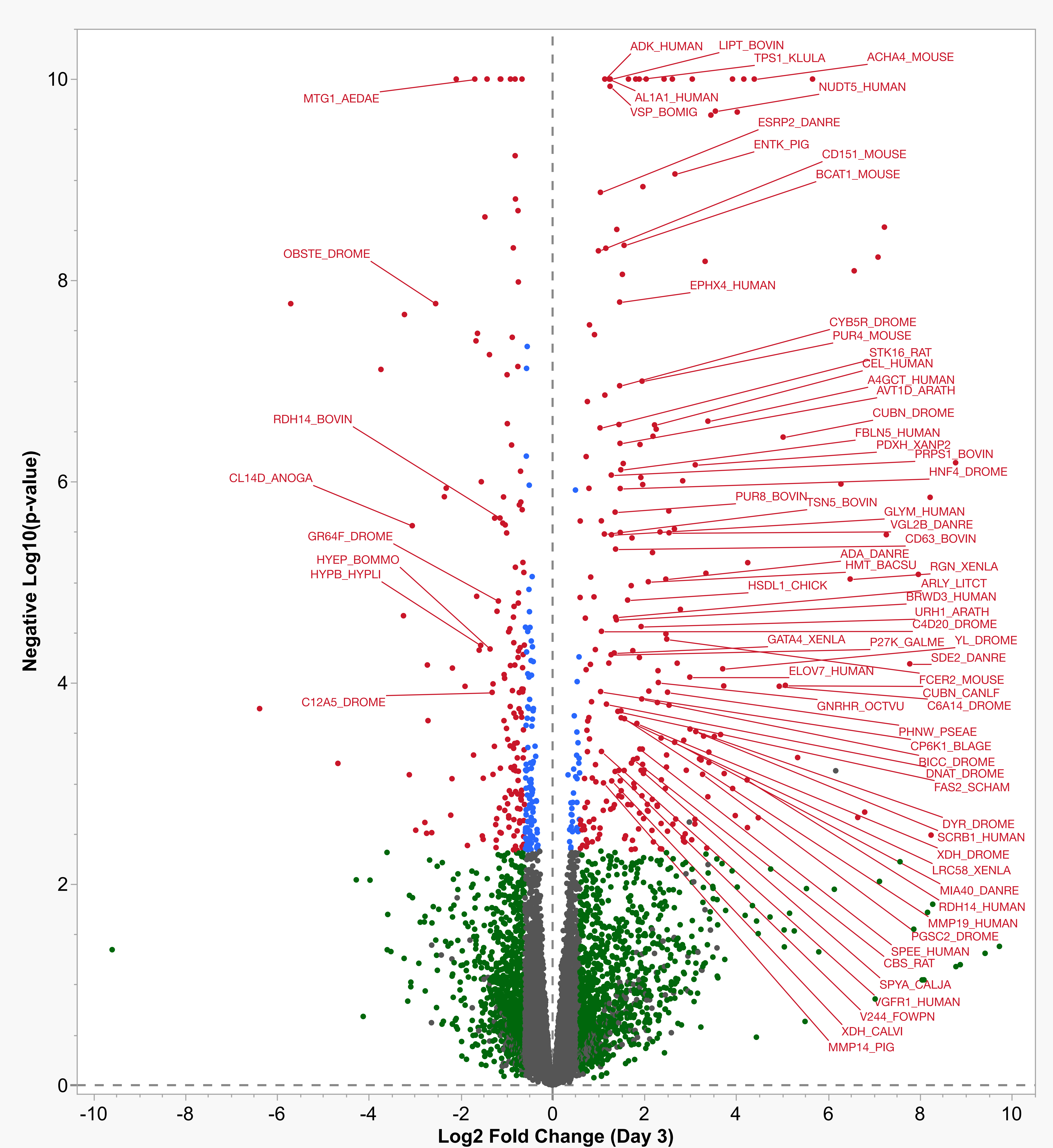

### Supplemental Materials S6

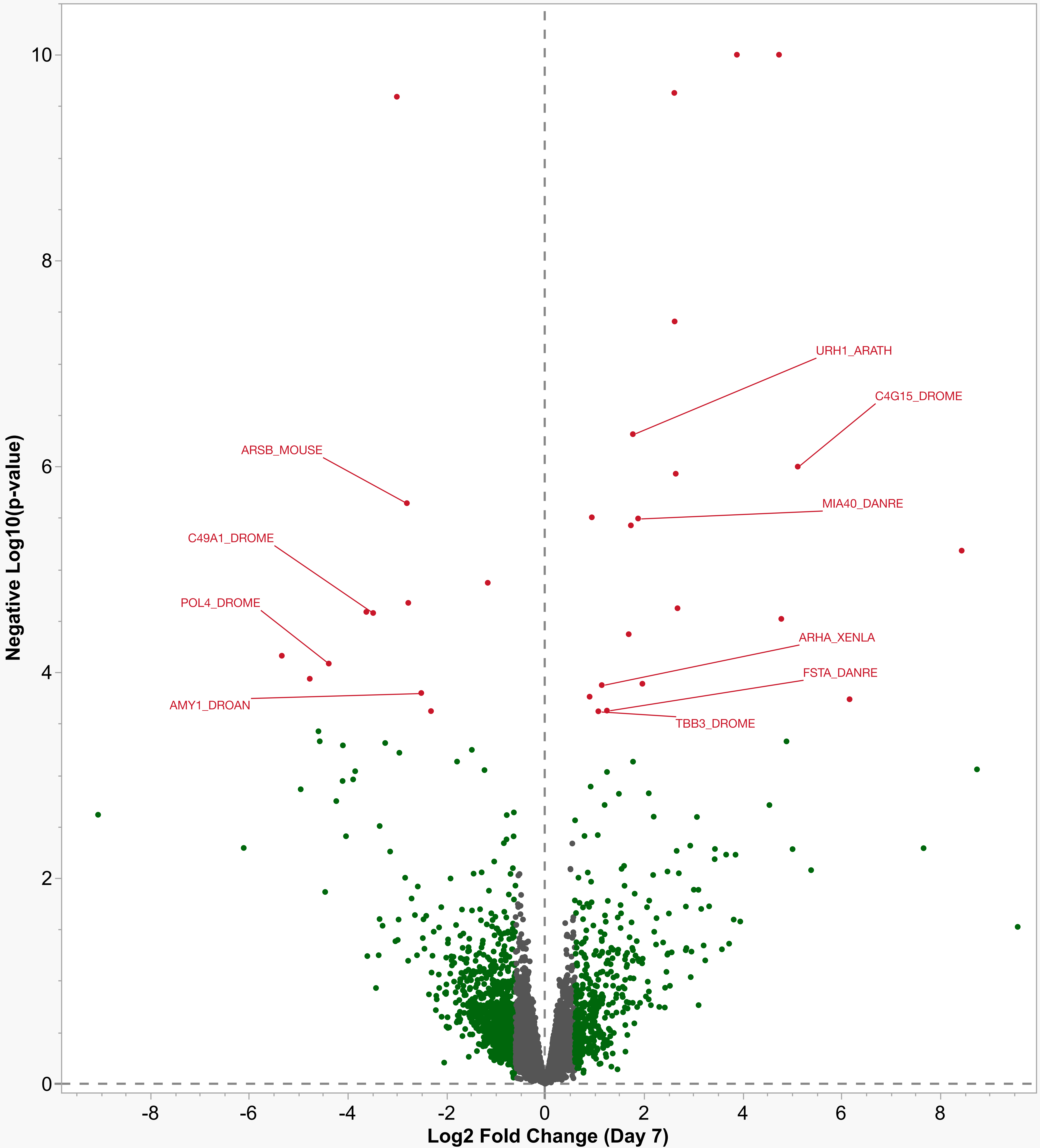
